# POU2AF2/OCA-T1 coactivates POU2F2 and defines a lineage-specific dependency in diffuse large B-cell lymphoma

**DOI:** 10.64898/2026.08.30.748132

**Authors:** Rima Tulaiha, Liam Shanley, Amanda Luvisotto, Ping Wang, Vipul Shukla, Zibo Zhao, Feng Yue, Ali Shilatifard, Christopher R Vakoc, Lu Wang

## Abstract

Lineage-restricted transcriptional programs establish cell identity and can create selective dependencies in cancer. Here, we identify POU2AF2, encoding the transcriptional co-activator OCA-T1, as a critical lineage-specific dependency in a subset of diffuse large B-cell lymphoma (DLBCL). Pan-cancer dependency analyses and patient cohorts reveal elevated POU2AF2 expression in genetically aggressive DLBCL, where its depletion markedly suppresses tumor growth *in vitro* and *in vivo*. Mechanistically, POU2AF2 cooperates with the B-cell lineage– defining transcription factor POU2F2 (OCT2) to activate lymphocyte activation gene programs through direct chromatin engagement, thereby sustaining malignant transcriptional networks. We further identified a key epigenetic regulatory axis composed of the lineage-specific transcription factor TCF3 and the histone methyltransferase SET1A-COMPASS that drives POU2AF2 expression downstream of B-cell receptor signaling. Single-cell transcriptomic analysis reveals that POU2AF2 marks and sustains an innate-like B1 B-cell population *in vivo*, a candidate cell of origin for lymphoma. Together, these findings define a lineage-restricted POU2AF2/POU2F2 transcriptional module, controlled by a TCF3/SET1A epigenetic network, that sustains both innate-like B-cell identity and malignant fitness in DLBCL. Our study uncovers a previously unrecognized lineage-specific transcriptional dependency and highlights POU2AF2 and its associated regulatory circuitry as potential therapeutic targets in aggressive B-cell malignancies.

## Introduction

Non-Hodgkin Lymphoma (NHL) accounts for 4% of all cancers in the United States, and it is estimated that 19,390 deaths from this disease will occur in the United States in 2025(*1*). NHL is a general category of lymphoma that begins in the lymphatic system, part of the body’s germ-fighting immune system(*1*). There are many subtypes that fall under the NHL category, and the classification of lymphoma depends on what type of lymphocyte is affected (B-cells or T-cells), how mature the cells are when they become cancerous, and other factors(*2*). B-cell lymphoma is a major form of NHL that originates in B-cell lymphocytes, which are responsible for producing antibodies in the immune system to combat infections(*3, 4*). The prevailing subtype of B-cell lymphoma is diffuse large B-cell lymphoma (DLBCL), a particularly aggressive subtype that continues to be the most common lymphoid malignancy in adults.

Various genetic and molecular alterations can drive the development of B-cell lymphoma, and different subtypes of B-cell lymphoma have distinct genetic drivers. Several genetic alterations have been identified to play a role in the development and progression of B-cell lymphoma, such as BCL2/BCL6 rearrangement(*5, 6*) and loss-of-function mutations within tumor suppressor genes like p53(*7*). The current clinical therapy for B-cell lymphoma typically involves a combination of chemotherapy, immunotherapy, targeted therapies, or stem cell transplantation(*8*).

Emerging studies from various research groups have suggested that epigenetic plasticity and reprogramming play a central role in guiding the intricate and dynamic phenotypic shifts observed in B-cells during both immune response and lymphomagenesis(*9, 10*). Research has demonstrated that disruptions in the B-cell lymphoma epigenome and the biological consequences of changes in epigenetic factors contribute to lymphoma progression. Examples of these contributions include gain-of-function mutations in EZH2(*11, 12*), loss-of-function mutations in MLL4/*KMT2D*(*13, 14*), frequent genetic abnormalities affecting histone acetyltransferases CBP and p300(*15*), and abnormal DNA methylation patterns(*16*). Collectively, epigenetic lesions hold promise as biomarkers to enhance the accuracy of diagnosis and prognosis in B-cell lymphoma(*9*). Alongside conventional chemotherapy and immunotherapy approaches, epigenetic therapies are gaining momentum in light of such findings. Further research is necessary to fully comprehend the impact of the newly identified epigenetic lesions on the genesis of B-cell lymphoma, as they have the potential to provide significant insights and advance the development of improved biomarkers and therapies(*17*).

The POU2F family of transcription factors, which includes POU2F1 (OCT1), POU2F2 (OCT2), and POU2F3 (OCT11), are characterized by a conserved POU DNA-binding domain that enables them to modulate the transcription of genes involved in differentiation, proliferation, and immune responses(*18, 19*). Compared with the broadly expressed POU2F1, POU2F2 and POU2F3 exhibit more cell-type specificity. For instance, POU2F3 is a master regulator of tuft cells (chemosensory epithelial cells located in intestine, airways, and urethra) and tuft cell-like tumors such as the small cell lung cancer P-subtype(*20–22*). In POU2F2 conditional knockout mice, the germinal center B cells are formed normally; however, their proliferation is reduced and *in vivo* differentiation to antibody-secreting plasma cells is blocked(*23, 24*). It has been demonstrated that in DLBCL cells, POU2F2 is required for the proliferation of the tumor cells by regulating the expression of numerous essential genes including *STAT3*, *IL-10*, *ELL2*, *XBP1*, *MYC*, *TERT*, and *ADA*(*24*). In addition, the POU2F transcription factors often coordinate their activities via interactions with co-activators, co-repressors or act in a complex with transcription factors from the same or other families(*25*). POU2AF1, also known as BOB1 or OCA-B, has been identified as the first co-activator of POU2F2, and the expression of which is largely restricted to B-cell lineage(*26–28*). Additionally, POU2AF1 is critical for tumor cell viability and has been validated as a cancer dependent factor in DLBCL(*29, 30*).

We and our colleagues initially identified a previously uncharacterized co-activator of the POU2F family, termed OCA-T1, encoded by the gene which we subsequently designated as *POU2AF2*, in the context of tuft cells and tuft cell-like malignancies(*20, 31, 32*). Genetic depletion of POU2AF2 led to a complete loss of tuft cells in the trachea and small intestine of mice(*20*). Similarly, in tuft cell-like tumors such as SCLC-P, POU2AF2 is essential for cell viability by facilitating POU2F3 function at chromatin and maintaining the expression of cell identity genes(*20, 21*). However, as a newly characterized gene, whether *POU2AF2* is expressed and plays a role in other cancer types, and whether it can activate transcription factors other than POU2F3 remains to be determined. In this study, we aim to define a distinct role for POU2AF2 in both human and murine B-cell lymphoma and to elucidate how POU2AF2 integrates into transcriptional networks that sustain lymphoma maintenance and normal B-cell lineage viability *in vitro* and *in vivo*.

## Results

### POU2AF2/OCA-T1 is a lineage-specific dependent factor in a subset of diffuse large B-cell lymphoma

Our previous work identified POU2AF2 as one of the most essential factors in the P-subtype of small cell lung cancer(*20, 31, 32*). To assess whether this dependency extends beyond SCLC-P, we performed a pan-cancer gene dependency analysis across diverse tumor lineages (Fig. 1A). Interestingly, in addition to SCLC-P cells, we observed a distinct subset of tumors of lymphoid origin that exhibited pronounced sensitivity to POU2AF2 depletion (Fig. 1A). Tumors of lymphoid origin arise from immune cells, including B cells, T cells, and natural killer (NK) cells, and encompass major malignancies such as non-Hodgkin lymphoma, Hodgkin lymphoma, leukemia, and plasma cell neoplasms(*33*). To determine which lymphoid subtypes express POUAF2, we employed the GENT2 database(*34*) and analyzed gene expression profiles from 5,010 patients with hematologic malignancies. This analysis revealed that POU2AF2 is highly expressed in a distinct subset of diffuse large B-cell lymphoma (DLBCL) patient samples (Fig. 1B). Gene expression analysis using UCSC Xena(*35*) reveals that POU2AF2 is significantly upregulated in DLBCL patient samples relative to normal B cells (Fig. 1C), suggesting a potential oncogenic role for POU2AF2 in DLBCL pathogenesis. Additionally, POU2AF2 expression was examined in a TCGA cohort of 481 diffuse large B-cell lymphoma (DLBCL) patient samples(*36, 37*). Comparative analysis revealed 461 genes significantly up-regulated in the top 10% of POU2AF2-expressing tumors (Fig. S1A). Notably, these genes were enriched for the Stromal-1 transcriptional signature(*38*), reflecting pathways associated with extracellular matrix deposition and cell–cell interactions (Fig. 1D–F).

**Figure 1.**
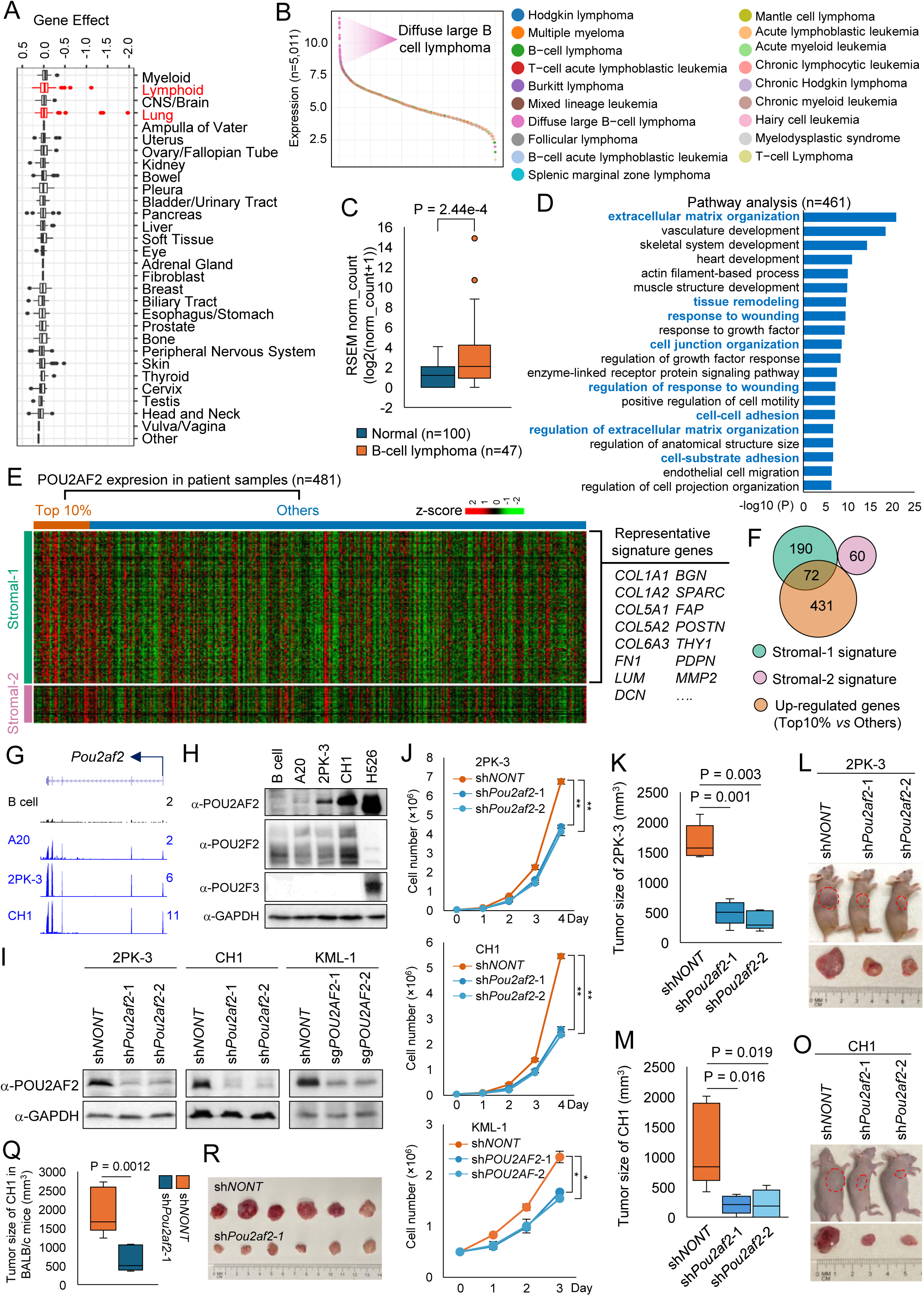
POU2AF2/OCA-T1 is a lineage-specific dependency in a subset of diffuse large B-cell lymphoma. A) Gene dependency scores for *POU2AF2* across diverse tumor lineages were obtained from the DepMap database. B) Dot plot shows *POU2AF2* gene expression across 5,010 blood cancer patient samples. C) The expression levels of *POU2AF2* in normal B cells and B cell lymphoma were retrieved from UCSC Xena database(*35*). The box plot compares *POU2AF2* gene expression between normal samples and B cell lymphoma. D) A cohort of 481 DLBCL patient samples was stratified into two groups based on POU2AF2 expression levels (top 10% vs. remaining 90%). Differential expression analysis identified 461 genes significantly upregulated in POU2AF2-high tumors. These genes were subjected to pathway enrichment analysis using Metascape. Enrichment significance is presented as −log₁₀(P), as calculated by Metascape (v3.5). E) Stromal-1 and Stromal-2 signature genes were retrieved from previous studies(*38*). The heatmap shows the expression levels of Stromal-1 and Stromal-2 signature genes between the two groups. F) The Venn diagram shows the overlap between genes up-regulated in POU2AF2-high group samples and Stromal-1 or Stromal-2 signature. G) Representative tracks show *Pou2af2* mRNA levels in mouse splenic B cells and the B cell lymphoma cell lines A20, 2PK-3, and CH1. H) Protein levels of POU2AF2, POU2F2, and POU2F3 in normal splenic B cells and B cell lymphoma cell lines (A20, 2PK-3, and CH1) were determined by western blot. The SCLC cell line NCI-H526 was used as a positive control for POU2AF2 expression, n = 2. I-J) Three different B-cell lymphoma cell lines, 2PK-3, CH1, and KML-1 were transduced with nontargeting or two independent *Pou2af2* shRNAs or sgRNAs. The protein levels of POU2AF2 were determined by western blot, n=2 (I), and the cell growth ability for each cell was determined by cell counting assay, n=3 (J). Data are represented as mean ± SD. A two-tailed unpaired Student’s t-test was used for statistical analysis. **P < 0.01; *P < 0.05. K-L) A total of 1×10⁶ 2PK-3 cells transduced with nontargeting or two independent *Pou2af2* shRNAs were inoculated into the right flank of athymic nude mice (n = 5 per group). Tumor volumes were measured at the experimental endpoint (K), with representative tumors shown (L). M-O) A total of 1×10^6^ CH1 cells transduced with nontargeting or two independent *Pou2af2* shRNAs were inoculated into the right flank of athymic nude mice (n = 5 per group). Tumor volumes (M) and representative tumors (O) are shown. Q-R) A total of 1×10^6^ CH1 cells transduced with nontargeting or *Pou2af2* shRNA were inoculated into BALB/c mice (n = 6 per group). Tumor volumes (Q) and representative tumors (R) are shown.

Next, we examined a panel of human and mouse B-cell lymphoma cell lines which lack typical tuft cell gene signature (Fig. S1A, B) to determine POU2AF2 protein levels. Consistent with that we observed in patient samples, high levels of POU2AF2 expression were detected in both human and mouse DLBCL cell lines, whereas POU2AF2 expression was undetectable in normal B cells isolated from mouse spleen (Fig. 1G, H, S1C). Notably, as cells of the B-cell lineage, the B-cell lymphoma cell lines expressed high levels of POU2F2, rather than POU2F3 (Fig. 1H). We then depleted POU2AF2 by two distinct shRNAs in three different B-cell lymphoma cell lines (Fig. S1D-F, 1I) to determine whether POU2AF2 represents a functional dependency required for lymphoma cell viability. As a result, POU2AF2 depletion markedly reduced tumor cell growth *in vitro* across all three cell lines we tested (Fig. J). To evaluate this requirement *in vivo*, we inoculated POU2AF2 wild-type or depleted cells into nude mice and observed a pronounced reduction in tumor growth upon POU2AF2 loss in both 2PK-3 cells (Fig. 1K, L) and CH1 cells (Fig. 1M, O). Consistently, POU2AF2 depletion also significantly suppressed CH1 tumor growth in immunocompetent BALB/c mice (Fig. 1Q, R). Together, these findings identify B-cell lymphoma as an additional POU2AF2 -dependent cancer type and demonstrate that POU2AF2 expression is required for robust B-cell lymphoma growth both *in vitro* and *in vivo*.

### POU2AF2/OCA-T1 controls the expression of lymphocyte activation genes at chromatin

To define how POU2AF2 regulates gene expression and to delineate the transcriptional program controlled by this cell-type-specific factor, we depleted POU2AF2 using two independent shRNAs in the CH1 cell line, which expresses the highest POU2AF2 level among all the cell lines tested, followed by RNA-seq analysis (Fig. 2A). Notably, pathway analysis of differentially expressed genes identified lymphocyte activation programming (*e.g. Ms4a1*, *Bcl11a*, *Zbtb32,* etc.) were substantially enriched among the downregulated genes upon POU2AF2 depletion (Fig. 2B). Consistent with our observations in CH1 cells, depletion of POU2AF2 in 2PK-3 cells also resulted in transcriptional repression of lymphocyte activation genes (Fig. S2A, B). To investigate the mechanism by which POU2AF2 regulates gene expression in B-cell lymphoma cells, we first determined the subcellular distribution of POU2AF2 protein. The subcellular fractionation analysis revealed that vast majority of POU2AF2 proteins were associated with chromatin, specifically within the insoluble nuclear fraction (Fig. 2C), suggesting that POU2AF2 may function as a transcriptional co-activator in B-cell lymphoma cells, analogous to its role in tuft cell lineages. Indeed, the ChIP-seq analysis identified 12,937 high-confidence POU2AF2 binding sites in CH1 cells (Fig. 2D, S2C). Interestingly, only approximately one-third of these sites were enriched at active enhancer regions (Fig. 2E, Cluster 1 peaks), whereas more than two-thirds are localized to promoter regions (Fig. 2E, Cluster 2 peaks). This chromatin-binding pattern of POU2AF2 in B-cell lymphoma cells contrasts distinctly with that observed in SCLC-P, in which POU2AF2 predominantly occupies distal enhancer elements (*31, 39*). These findings indicate that POU2AF2 may engage distinct regulatory mechanisms in B-cell lymphoma cells.

**Figure 2.**
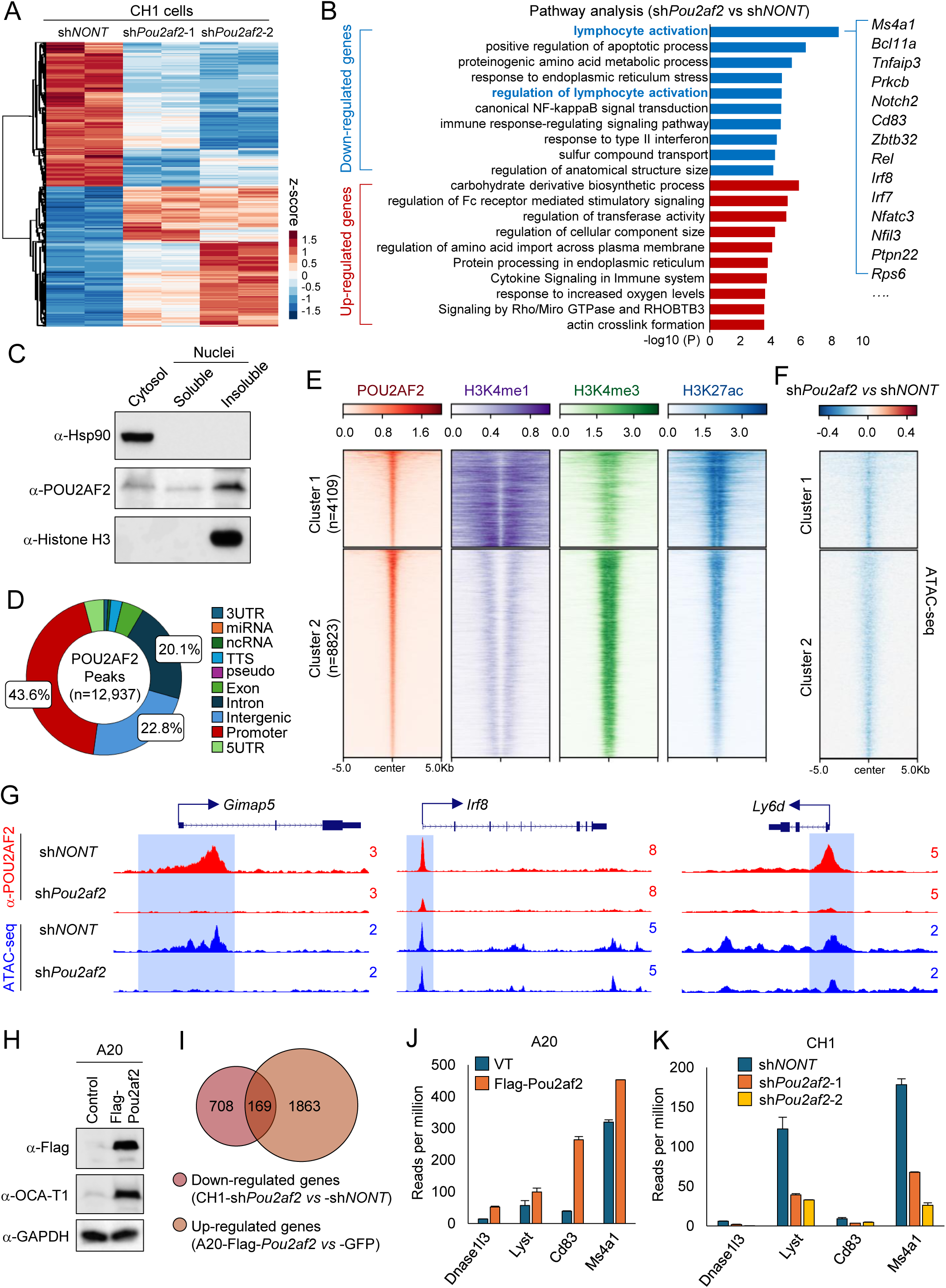
POU2AF2/OCA-T1 controls the expression of lymphocyte activation genes at chromatin. A) RNA-seq was performed in CH1 cells transduced with nontargeting or *Pou2af2*-specific shRNAs for 4 days. Heatmap shows the differentially expressed genes (n = 2). B) Metascape pathway enrichment analysis of genes downregulated or upregulated upon POU2AF2 depletion. Enrichment significance is presented as −log10(P), calculated using Metascape v3.5. C) Subcellular fractionation followed by western blot analysis of POU2AF2 protein levels in cytosolic, soluble nuclear, and insoluble nuclear fractions. HSP90 and histone H3 were used as cytoplasmic and insoluble nuclear controls, respectively, n=2. D) Pie chart shows the genomic annotation and distribution of POU2AF2 peaks. E) POU2AF2 peaks were grouped into two clusters by k-means clustering, and histone modification signals were centered on POU2AF2 peaks within each cluster. F) ATAC-seq was performed in CH1 cells transduced with nontargeting or *Pou2af2*-specific sgRNAs for 4 days. Log2 fold changes in chromatin accessibility heatmap was shown when centered on POU2AF2 peaks as ranked in (E). G) Representative tracks illustrate changes in chromatin accessibility at lymphocyte activation-associated genes upon POU2AF2 depletion. H) A20 cells were transduced with lentivirus expressing empty vector or FLAG-tagged POU2AF2, and POU2AF2 protein levels were determined by western blot, n=2. I) Venn diagram analysis shows the overlap between genes downregulated upon POU2AF2 depletion in CH1 cells and genes upregulated following POU2AF2 overexpression in A20 cells. J) Bar plots shows the expression of *Dnase1l3*, *Lyst*, *Cd83*, and *Ms4a1* in A20 cells expressing empty vector or FLAG-tagged POU2AF2, n = 2. K) Bar plot shows the expression of *Dnase1l3*, *Lyst*, *Cd83*, and *Ms4a1* in CH1 cells transduced with nontargeting or *Pou2af2*-specific sgRNAs, n = 2.

To further investigate how POU2AF2 regulates chromatin states and accessibility, we performed ATAC-seq analysis in CH1 wild-type and POU2AF2 depleted cells (Fig. S2D). Loss of POU2AF2 resulted in a marked decrease in chromatin accessibility at POU2AF2 occupied loci (Fig. 2F). Notably, most of the reduced ATAC-seq peaks were located at chromatin regions of genes involved in immune responses and lymphocyte activation (Fig. 2G, S2E). To determine whether POU2AF2 overexpression could restore the expression of these genes, we ectopically expressed POU2AF2 in A20 cells, which normally express low POU2AF2 levels (Fig. 2H). Among the genes significantly downregulated upon POU2AF2 depletion in CH1 cells, 169 were significantly induced upon POU2AF2 overexpression in A20 cells based on RNA-seq analysis (Fig. 2I). Pathway analysis revealed that lymphocyte activation was the most significantly enriched pathway among the up-regulated genes (Fig. 2J, K, S2F). Collectively, these results reveal that POU2AF2 functions as a major transcriptional activator, specifically regulating lymphocyte activation genes in B-cell lymphoma cells.

### POU2AF2/OCA-T1 co-activates POU2F2 in B cell lymphoma

In tuft cells and tuft cell-like tumor cells, POU2AF2 forms a heterodimer with the master transcription factor POU2F3 and functions as a transcriptional co-activator at enhancer regions(*31, 40*). Within the B-cell lineage, POU2F2 was predominantly expressed, whereas POU2F3 expression was minimal (Fig. 1E). Therefore, to investigate how POU2AF2 regulates transcriptional programs through its co-activator activity, we performed integrated motif analyses of POU2AF2 ChIP-seq (Fig. 2E) and ATAC-seq datasets generated from POU2AF2 wild type and depleted cells (Fig. S2D). Interestingly, HOMER motif analysis revealed that motifs corresponding to POU2F2 and POU2F3 were identified as the top two enriched hits (Fig. 3A). Given that B-cell lymphoma cells express minimal levels of POU2F3 but high levels of POU2F2 (Fig. 1E), we hypothesized that POU2AF2 may function as a transcriptional co-activator of POU2F2 in this cellular context. Supporting this model, co-immunoprecipitation experiments revealed a robust protein-protein interaction between POU2AF2 and POU2F2 in different cell lines, including CH1 (Fig. 3B), 2PK-3 (Fig. S3A), and KML-1 cells (Fig. S3B). Additionally, size-exclusion chromatography of nuclear extracts from CH1 cells revealed a prominent co-elution of POU2F2 with POU2AF2, in contrast to POU2AF1, the known B-cell specific co-activator of POU2F2(*27, 41*). These data indicate a potential dominant role for the POU2F2/POU2AF2 heterodimer in B-cell lymphoma. Consistently, ChIP-seq analysis revealed extensive genomic co-occupancy of POU2AF2 and POU2F2 in CH1 cells (Fig. 3D-G). Additionally, similar results were obtained in human KML-1 cell line, in which a strong co-localization between POU2AF2 and POU2F2 was also confirmed by ChIP-seq analysis at genome wide levels (Fig. S3C-F).

**Figure 3.**
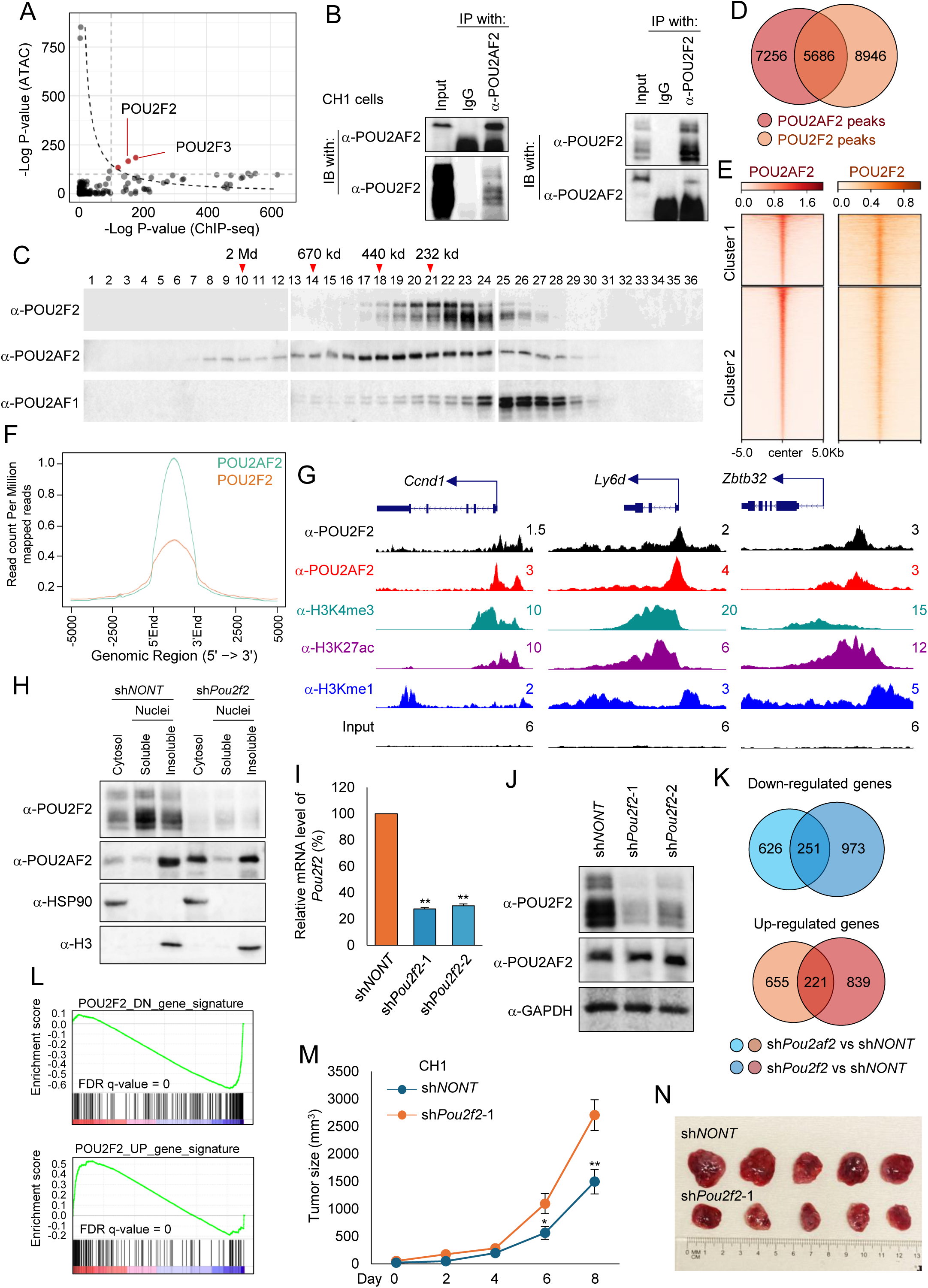
POU2AF2/OCA-T1 co-activates POU2F2 in B cell lymphoma. A) HOMER motif analysis of POU2AF2 ChIP-seq peaks and ATAC-seq regions with reduced accessibility upon POU2AF2 depletion. Negative log p-values for motif enrichment are shown for ChIP-seq (x-axis) and ATAC-seq (y-axis), revealing significant enrichment of POU2F2 and POU2F3 motifs in both. B) Protein-protein interaction between POU2AF2 and POU2F2 in CH1 cells was detected by immunoprecipitation, n=2. C) Nuclear extracts from CH1 cells were fractionated by size-exclusion chromatography, and POU2F2, POU2AF2, and OCA-B protein levels in eluted fractions were analyzed by western blot, n=2. D) POU2F2 ChIP-seq in CH1 cells. Venn diagram shows the overlap between POU2AF2 and POU2F2 binding peaks. E) POU2F2 ChIP-seq signal in CH1 cells were centered on the two clusters of POU2AF2 peaks in Fig. 2E. F) Genome-wide average plot shows the co-occupancy of POU2F2 and POU2AF2. G) Representative tracks illustrating co-localization of POU2F2 and POU2AF2 at promoter regions of lymphocyte activation genes. H) CH1 cells were transduced with nontargeting or *Pou2f2*-specific shRNA for 96 h, followed by subcellular fractionation. POU2F2 and POU2AF2 protein levels in each fraction were determined by western blot. HSP90 and histone H3 were used as cytoplasmic and insoluble nuclear controls, respectively, n = 2). I-J) CH1 cells were transduced with nontargeting or two independent *Pou2f2*-specific shRNAs for 96 h, and *Pou2f2* mRNA (I) and protein (J) levels were assessed by RT-qPCR (n=3) and western blot (n=2), respectively. K-L) RNA-seq analysis of CH1 cells transduced with nontargeting or *Pou2f2*-specific shRNA #1 for 96 h. Venn-diagram (K) and GSEA (L) analysis show the overlap between up- and downregulated genes in POU2F2- and POU2AF2-depleted cells. M-N) A total of 1×10^6^ CH1 cells transduced with nontargeting or *Pou2f2*-specific shRNA were inoculated into the right flank of athymic nude mice (n=5 per group). Tumor growth was monitored every 2 days starting 2 weeks after inoculation (M), and representative tumors are shown (N). Data are represented as mean ± SEM. A two-tailed unpaired Student’s t-test was used for statistical analysis. **P < 0.01; *P < 0.05.

As a transcriptional co-activator, POU2AF2 lacks a recognizable DNA-binding motif and is instead recruited to chromatin through interaction with transcription factors via its N-terminal peptide(*42*). Accordingly, depletion of partner transcription factors POU2F3 results in a marked loss of POU2AF2 chromatin association in SCLC-P cells(*31*) . To determine whether POU2F2 similarly mediates POU2AF2 recruitment to chromatin in B-cell lymphoma, we depleted POU2F2 in CH1 cells and examined POU2AF2 protein distribution across cellular fractions. As a result, POU2F2 depletion led to a dramatic reduction of chromatin-bound POU2AF2, accompanied by its accumulation in the cytoplasmic fraction (Fig. 3H). These findings indicate that POU2AF2 is recruited to chromatin through a conserved transcription factor-dependent mechanism in both tuft-cell and B-cell lineages.

Finally, to define the functional interplay between POU2F2 and POU2AF2 in the regulation of gene expression in B-cell lymphoma, we depleted POU2F2 using two independent shRNAs in CH1 cells (Fig. 3I, J) and performed RNA-seq analysis. Venn diagram analysis (Fig. 3K) and GSEA analysis (Fig. 3L) revealed a substantial overlap between genes regulated by POU2F2 and POU2AF2 in CH1 cells, including genes involved in lymphocyte activation pathway (Fig. S3G). POU2F2 is a B-cell-restricted transcription factor(*43*) that is indispensable for B-cell development, proliferation, and function(*44, 45*). Its expression is markedly elevated in B-cell lymphoma (Fig. S3H) and inversely correlated with patient survival (Fig. S3I). Similar to POU2AF2, DepMap analysis indicates that POU2F2 is selectively required for the viability of tumors of lymphoid origin (Fig. S3J). Consistent with these clinical observations, loss of POU2F2 significantly impaired tumor growth *in vivo* (Fig. 3M, N). Together, these findings establish a critical role for the POU2AF2/POU2F2 heterodimer in coordinating transcriptional programs and driving tumor growth in B-cell lymphoma.

### TCF3 links BCR signaling to *POU2AF2* activation and lymphoma progression

To investigate the mechanisms underlying transcriptional activation of POU2AF2 in B-cell lymphoma, we compared gene expression profiles across three mouse B-cell lymphoma cell lines, A20, 2PK-3 and CH1. Among these, CH1 cells express the highest POU2AF2 levels (Fig. 1G, H) and display markedly accelerated growth *in vivo* relative to A20 and 2PK-3 cells (Fig. 4A). Venn diagram analysis identified 1,567 genes that were significantly upregulated in CH1 cells compared with both A20 and 2PK-3 cells (Fig. 4B), including *POU2AF2* gene (Fig. S4A, B). Pathway enrichment analysis of these genes revealed lymphocyte activation as the top-ranked pathway (Fig. 4C), consistent with the POU2AF2 targeting pathways (Fig. 2B, Fig. S2B, S2F), and the pronounced cell aggregate formation observed during CH1 cell culture (Fig. 4D). Indeed, pharmacological inhibition of B-cell receptor (BCR) signaling with either R406 or ibrutinib markedly reduced POU2AF2 expression at both the mRNA (Fig. 4E) and protein levels (Fig. 4F). RNA-seq analysis further revealed that 94 POU2AF2 target genes associated with immune responses and lymphocyte activation were concomitantly downregulated upon BCR inhibition (Fig. 4G, H).

**Figure 4.**
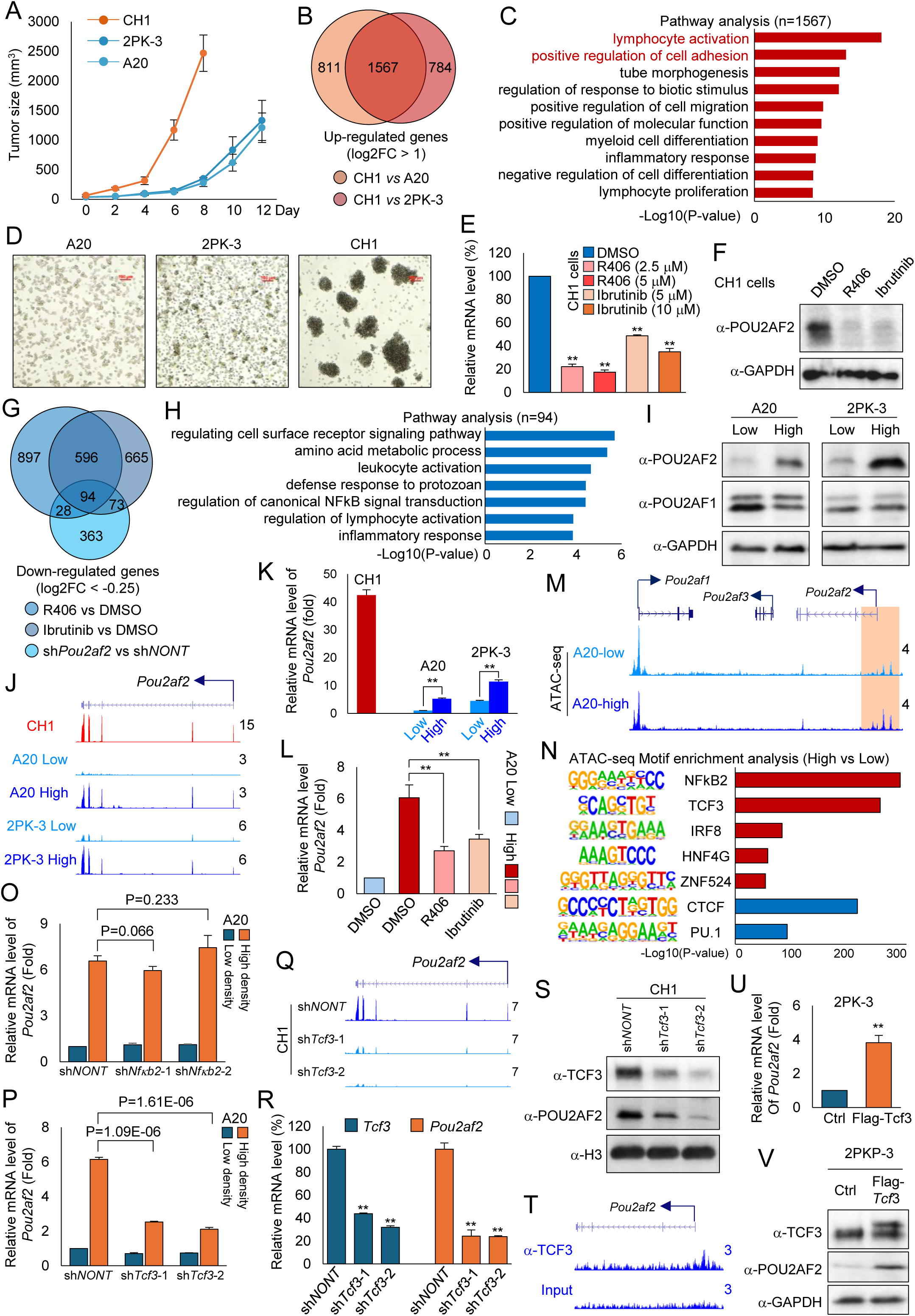
TCF3 links BCR signaling to *POU2AF2* activation and B-cell lymphoma progression. A) A total of 1×10^6^ CH1, 2PK-3, or A20 cells were inoculated into the right flank of athymic nude mice (n=5 per group). Tumor growth was measured every 2 days starting 2 weeks after inoculation. B) Venn diagram shows the genes commonly upregulated in CH1 cells compared with A20 or 2PK-3 cells. C) Metascape pathway enrichment analysis of genes upregulated in CH1 cells identified in (B). Enrichment significance is shown as −log_10_(P), calculated using Metascape v3.5. D) Representative images show cell morphology of A20, 2PK-3, and CH1 cells. E) CH1 cells were treated with various concentrations of R406 or ibrutinib for 24 h, and *Pou2af2* mRNA levels were quantified by RT-qPCR, n = 3. Data are represented as mean ± SD. A two-tailed unpaired Student’s t-test was used for statistical analysis. **P < 0.01; *P < 0.05. F) CH1 cells were treated with 5 μM R406 or ibrutinib for 24 h, and POU2AF2 protein levels were determined by western blot, n=2. G) Venn diagram analysis shows the overlap between genes downregulated in CH1 cells treated with R406 or ibrutinib and genes downregulated upon POU2AF2 depletion. H) Metascape pathway enrichment analysis of the 94 overlapping genes identified in (G). I) A20 and 2PK-3 cells were cultured at low (0.5 × 10⁵ cells/ml) or high (2.5 × 10⁵ cells/ml) density for 96 h. POU2AF2 and POU2AF1 protein levels were determined by western blot. GAPDH was used as internal control, n = 2. J) Representative tracks show *Pou2af2* gene expression in CH1, A20, and 2PK-3 cells cultured under low- or high-density conditions. K) RT-qPCR analysis of *Pou2af2* mRNA levels in A20 and 2PK-3 cells cultured under low- or high-density conditions compared to that in CH1 cells, n=3. Data are represented as mean ± SD. A two-tailed unpaired Student’s t-test was used for statistical analysis. **P < 0.01; *P < 0.05. L) RT-qPCR analysis of *Pou2af2* mRNA levels in CH1 cells and in A20 cells treated with DMSO, R406, or ibrutinib, n = 3. Data are represented as mean ± SD. A two-tailed unpaired Student’s t-test was used for statistical analysis. **P < 0.01; *P < 0.05. M-N) ATAC-seq analysis in A20 cells cultured under low- or high-density conditions. Representative tracks show chromatin accessibility at the *Pou2af2* gene locus (M), and motif analysis identifies enriched transcription factor motifs in regions with increased or decreased accessibility (N). O) A20 cells transduced with nontargeting or two independent *Nfκb2*-specific shRNAs were cultured under low- or high-density conditions for 4 days, and *Pou2af2* mRNA levels were measured by RT-qPCR, n = 3. Data are represented as mean ± SD. A two-tailed unpaired Student’s t-test was used for statistical analysis. P) A20 cells transduced with nontargeting or two independent *Tcf3*-specific shRNAs were cultured under low- or high-density conditions for 4 days, followed by RT-qPCR analysis of *Pou2af2* gene expression, n = 3. Data are represented as mean ± SD. A two-tailed unpaired Student’s t-test was used for statistical analysis. Q-R) CH1 cells transduced with nontargeting or *Tcf3*-specific shRNAs for 4 days were analyzed for *Pou2af2* gene expression by RNA-seq (n=2) (Q) and RT-qPCR (n=3). A two-tailed unpaired Student’s t-test was used for statistical analysis. Data are represented as mean ± SD. A two-tailed unpaired Student’s t-test was used for statistical analysis. **P < 0.01; *P < 0.05 (R). S) Western blot analysis of TCF3 and POU2AF2 protein levels in CH1 cells transduced with nontargeting or *Tcf3*-specific shRNAs, n = 2). T) ChIP-seq analysis in CH1 cells using a TCF3-specific antibody. Representative tracks show TCF3 occupancy at the *Pou2af2* promoter. U-V) 2PK-3 cells were transduced with lentivirus expressing empty vector or FLAG-tagged TCF3. *Pou2af2* mRNA (U) and protein (V) levels were determined by RT-qPCR (n=3) and western blot (n=2), respectively. Data are represented as mean ± SD. A two-tailed unpaired Student’s t-test was used for statistical analysis. **P < 0.01; *P < 0.05.

Interestingly, we found that when the POU2AF2 low-expressing cell lines A20 and 2PK-3 were cultured at high cell density to mimic the CH1 cells, the POU2AF2 expression levels were robustly induced at both the mRNA (Fig. 4I, J) and protein levels (Fig. 4K). Notably, this cell aggregation-induced POU2AF2 upregulation could also be suppressed by pharmacological BCR inhibition (Fig. 4L). We therefore exploited cell density as a perturbation to identify the transcription factors regulating POU2AF2 expression downstream of BCR signaling. ATAC-seq analysis in A20 cells revealed a substantial increase in chromatin accessibility at POU2AF2 promoter region (Fig. 4M) as well as at genome wide levels (Fig. S4C-E). Based on these results, we conducted global motif enrichment analysis to identify transcription factor motifs that were significantly altered between low- and high-density conditions. Among the top candidates, NFκB2 and TCF3 were selected for further validation (Fig. 4N). As a result, we found depletion of TCF3 (Fig. S4F), but not NFκB2 (Fig. S4G), significantly impaired density-induced activation of POU2AF2 expression in A20 cells (Fig. 4O, P). To validate this result in POU2AF2 highly expressed cells, we depleted TCF3 using two independent shRNAs in CH1 cells, in which BCR signaling is constitutively active and expresses high levels of TCF3 (Fig. S4h). We found that TCF3 depletion led to a substantial reduction in POU2AF2 expression at both the mRNA (Fig. 4Q, R) and protein levels (Fig. 4S), without affecting the expression of POU2AF1 (Fig. S4I), the predominant co-activator of POU2F2(*28*). To exclude potential off-target effects by the shRNAs, TCF3 was depleted by CRISPR in CH1-Cas9 cells, which similarly resulted in a marked reduction of POU2AF2 expression (Fig. S4J, K).

By ChIP-seq analysis, we have detected strong TCF3 occupancy at the *POU2AF2*gene promoter in CH1 cells, indicating a direct transcriptional regulation of POU2AF2 by TCF3 (Fig. 4T). Consistent with the results that we observed in CH1 cells, ectopic expression of TCF3 in the POU2AF2 low expressed cell line 2PK-3 was sufficient to induce POU2AF2 expression at both the mRNA (Fig. 4U, Fig. S4L) and protein levels (Fig. 4V). Similar to POU2AF2 depletion *in vivo*, loss of TCF3 markedly impaired tumor growth in animal models (Fig. S4M, N), in agreement with dependency analyses from the DepMap database (Fig. S4O). Collectively, these findings establish TCF3 as a key mediator of BCR-dependent POU2AF2 activation in B-cell lymphoma.

### SET1A COMPASS selectively controls *POU2AF2* expression in B-cell lymphoma

We have shown a transcriptional activation of POU2AF2 expression driven by the lineage-specific transcription factor TCF3 in multiple B-cell lymphoma cell lines. Interestingly, as we noticed in the A20 cell line, depletion of TCF3 did not completely abolish POU2AF2 activation under high-density culture conditions (Fig. 4P), suggesting the involvement of additional regulatory mechanisms. Therefore, to investigate whether epigenetic regulation also contributes to POU2AF2 expression in B-cell lymphoma cells, we compared chromatin states in A20 cells cultured under low- and high-density conditions by profiling active and repressive histone marks. As shown in Fig. 5A, we observed a substantial increase in the active histone mark H3K4me3 at specific genomic loci that exhibited transcriptional activation as determined by RNA-seq analysis (Fig. 5B, C). Pathway analysis revealed that these genes are significantly enriched in cell-activation related pathways (Fig. 5D). Among the genes associated with the Cluster 1 chromatin signature, the *Pou2af2* gene promoter displayed a pronounced increase in active histone marks, including H3K4me3 and H3K27ac (Fig. 5E). In contrast, no significant changes in either gene expression or histone modifications were detected at the *Pou2af1* gene promoters (Fig. S5A). Consistently, selective enrichment of H3K4me3 at the Pou2af2 promoter under high-density culture conditions was also observed in 2PK-3 cells (Fig. S5B, C), supporting the existence of a specific epigenetic mechanism governing *Pou2af2* expression. Moreover, these active histone marks were constitutively elevated at the *Pou2af2* promoter in CH1 cells (Fig. 5F), which express substantially higher levels of POU2AF2 than A20 or 2PK-3 cells (Fig. 1D).

**Figure 5.**
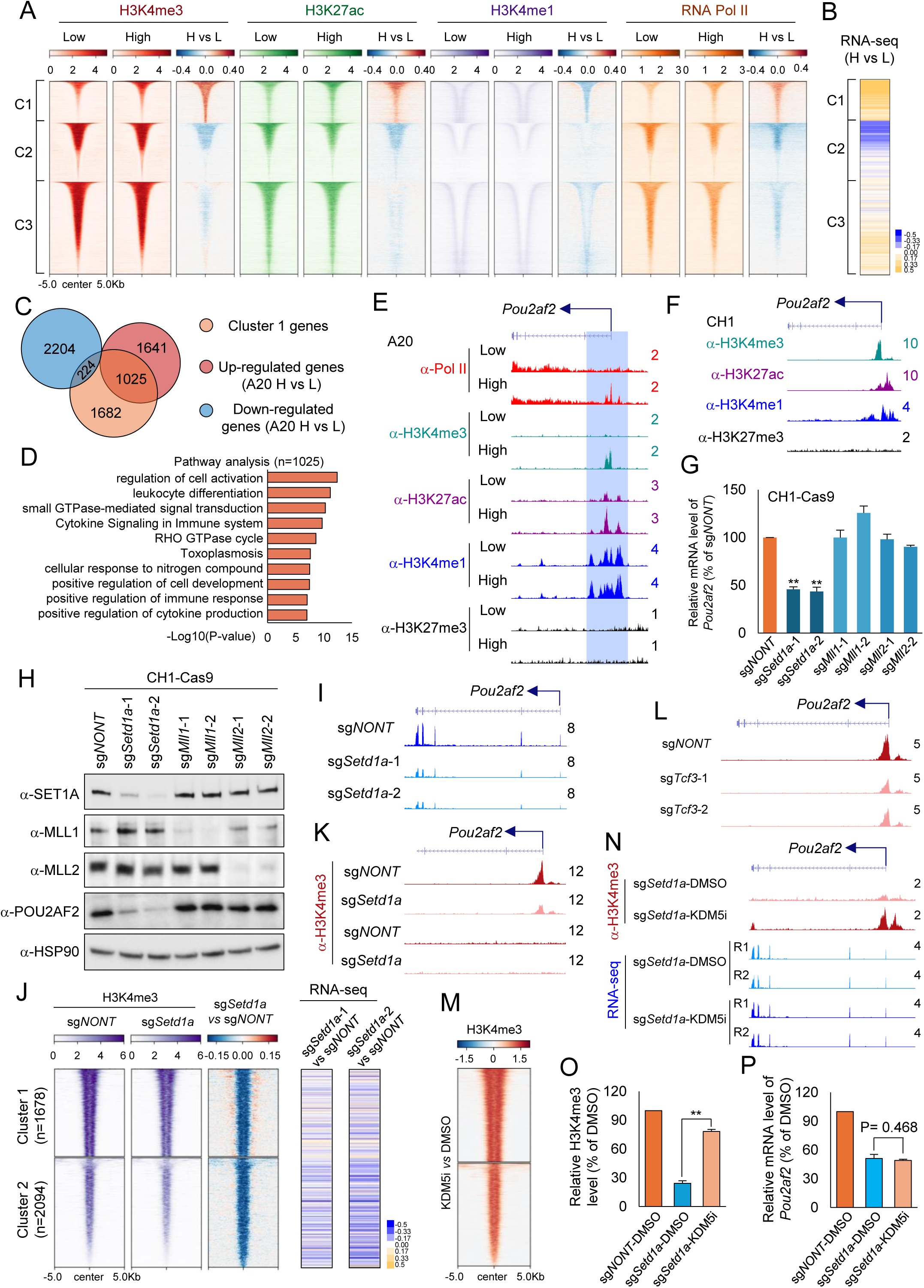
SET1A COMPASS selectively controls *POU2AF2* expression in B-cell lymphoma. A) ChIP-seq was performed to profile H3K4me1, H3K4me3, H3K27ac, and RNA polymerase II occupancy in A20 cells cultured under low- or high-density conditions. All peaks were grouped into three clusters by k-means clustering. Occupancy and log2FC heatmaps were shown. B) Heatmap analysis shows changes in expression of genes proximal to the corresponding ChIP-seq peaks defined in (A). C) Venn diagram shows the overlap of genes upregulated or downregulated under high-density culture conditions that are proximal to cluster 1 peaks defined in (A). D) Metascape pathway enrichment analysis of genes upregulated under high-density culture conditions and located near cluster 1 peaks. E) Representative tracks show Pol II, H3K4me3, H3K4me1, H3K27me3, and H3K27ac occupancy at the *Pou2af2* gene promoter in A20 cells cultured under low- or high-density conditions. F) Representative tracks show H3K4me3, H3K27ac, H3K27me3, and H3K4me1 occupancy at the *Pou2af2* gene promoter in CH1 cells. G-H) CH1 cells stably expressing Cas9 were transduced with nontargeting sgRNA or two independent sgRNAs targeting *SET1A*, *MLL1*, or *MLL2*. The *Pou2af2* mRNA in each group was determined by RT-PCR (n=3) (G), and the protein levels were determined by western blot (n=2) (H). Data are represented as mean ± SD. A two-tailed unpaired Student’s t-test was used for statistical analysis. **P < 0.01; *P < 0.05. I) Representative RNA-seq tracks show *Pou2af2* gene expression in CH1-Cas9 cells transduced with nontargeting sgRNA or *SET1A*-specific sgRNAs. J) ChIP-seq analysis of H3K4me3 in CH1-Cas9 cells transduced with nontargeting or SET1A-specific sgRNA. H3K4me3 peaks were divided into two clusters by k-means clustering (left). Log_2_ fold change heatmap shows a global reduction of H3K4me3 levels upon SET1A depletion (middle), and RNA-seq heatmap shows expression changes of genes proximal to the corresponding peaks (right). K) Representative tracks show H3K4me3 occupancy at the *Pou2af2* gene promoter in wild-type and SET1A-depleted CH1 cells. L) Representative tracks show H3K4me3 occupancy at the *Pou2af2* gene promoter in CH1-Cas9 cells transduced with nontargeting or *Tcf3*-specific sgRNAs. M) SET1A-depleted CH1 cells were treated with DMSO or 5 μM of the pan-KDM5 inhibitor KDM5C70 for 24 h, followed by H3K4me3 ChIP-seq. Log_2_ fold change heatmap shows a global increase in H3K4me3 levels upon KDM5 inhibition. N) Representative tracks show H3K4me3 occupancy at the *Pou2af2* gene promoter and *Pou2af2* gene expression in SET1A-depleted CH1 cells treated with DMSO or KDM5C70. O) The H3K4me3 occupancy at the *Pou2af2* gene promoter in wild-type and SET1A-depleted CH1 cells treated with DMSO or KDM5C70 was validated by ChIP-qPCR, n=3. Data are represented as mean ± SD. P) The *Pou2af2* mRNA levels in wild-type and SET1A-depleted CH1 cells treated with DMSO or KDM5C70 were determined by RT-qPCR, n=3. Data are represented as mean ± SD. A two-tailed unpaired Student’s t-test was used for statistical analysis. **P < 0.01; *P < 0.05.

It has been well established that the COMPASS complex serves as the main histone H3K4 methyltransferase from yeast to mammals(*46–48*). In particular, H3K4me3 deposition is primarily catalyzed by the MLL1, MLL2, and SET1A COMPASS complexes(*48*). To identify which COMPASS complex is responsible for H3K4me3 enrichment at the *Pou2af2* gene promoter, we individually depleted SET1A, MLL1, and MLL2 using CRISPR in CH1 cells. Notably, depletion of SET1A, but not MLL1 or MLL2, resulted in a pronounced reduction in both POU2AF2 mRNA and protein levels (Fig. 5G-I) in CH1 cells, which exhibit constitutively high H3K4me3 levels at the *Pou2af2* gene promoter. Consistent with these observations, SET1A depletion led to a global decrease in H3K4me3 levels across the genome (Fig. 5J, left panel), accompanied by reduced expression of nearby genes (Fig. 5J, right panel). Pathway analysis revealed that these downregulated genes were significantly enriched for immune response and lymphocyte activation pathways (Fig. S5D, E).

Notably, we detected a significant occupancy of SET1A at *Pou2af2* gene promoter region in CH1 cells (Fig. S5F), and loss of SET1A resulted in a substantial reduction of H3K4me3 level (Fig. 5K; Fig. S5G), without affecting that at the *Pou2af1* gene locus (Fig. S5H) or *Pou2af1* gene expression (Fig. S5I). Interestingly, depletion of TCF3 similarly reduced H3K4me3 levels at the *Pou2af2* gene promoter (Fig. 5L), but not at the *Pou2af1* gene promoter (Fig. S5J), suggesting a functional cooperation and genetic interaction between TCF3 and the SET1A COMPASS complex. Collectively, these data demonstrate that the SET1A COMPASS directly regulates POU2AF2 transcription in B-cell lymphoma cells, potentially through coordinated interactions with lineage-specific transcription factors.

Previous studies have shown that KDM5 demethylases mediate the dynamic turnover of H3K4me3 following COMPASS depletion, and that pan-KDM5 inhibitors such as KDM5-C70 can restore global H3K4me3 levels in this context(*49*). To determine whether the reduced H3K4me3 levels observed upon SET1A depletion is causative for POU2AF2 transcriptional downregulation, we treated SET1A-depleted CH1 cells with the KDM5 inhibitor KDM5-C70(*49*). As expected, KDM5-C70 treatment led to a robust increase in H3K4me3 levels both genome-wide (Fig. 5M, Fig. S5K) and at the *Pou2af2* gene promoter (Fig. 5N). However, despite efficient restoration of H3K4me3, KDM5 inhibition failed to rescue *Pou2af2* gene expression in SET1A-depleted cells (Fig. 5N). This finding was further validated by ChIP-qPCR analysis (Fig. 5O) and real time PCR (Fig. 5P). Together, these results are consistent with previous reports indicating that H3K4me3 enrichment is a consequence rather than a driver of transcriptional activation(*50*).

### POU2AF2/OCA-T1 defines and sustains a distinct B1-lineage B-cell population *in vivo*

Previous studies have demonstrated that POU2AF2 is selectively expressed in normal tuft cells as well as in tuft cell-like tumors, including the P-subtype of small cell lung cancer(*20*). Genetic depletion of POU2AF2 induces profound apoptosis in SCLC-P tumor cells and completely abolishes tumor growth *in vivo*(*31*). Consistently, *Pou2af2* null mice exhibit a near-complete loss of tuft cells in both the trachea and small intestine(*20*). On the basis of these observations, we hypothesized that only specific B cell sub-populations express *POU2AF2*, given that its protein levels were barely detectable in bulk B-cell populations (Fig. 1D). To test this hypothesis, we isolated splenic B cells from BALB/c mice and subjected them to single-cell RNA sequencing (scRNA-seq) analysis. UMAP analysis revealed multiple B-cell subsets defined by distinct transcriptional programs (Fig. 6A). As expected, the B-cell-specific transcription factor *Pou2f2* (Fig. 6B, left panel) and its known co-activator *Pou2af1* were broadly expressed across the majority of B-cell populations (Fig. 6B, middle panel). Strikingly, *Pou2af2* expression was strongly enriched in a single, discrete cluster, which was annotated as activated memory B cells (Fig. 6B, right panel). To investigate whether *Pou2af2* expression reflects a distinct B-cell state program, we performed pseudotemporal trajectory analysis, which revealed a continuous differentiation trajectory from immature to activated B-cell states. Within this trajectory, *Pou2af2* expressing B cells occupied the most differentiated state of splenic B cells (Fig. 6C, S6A).

**Figure 6.**
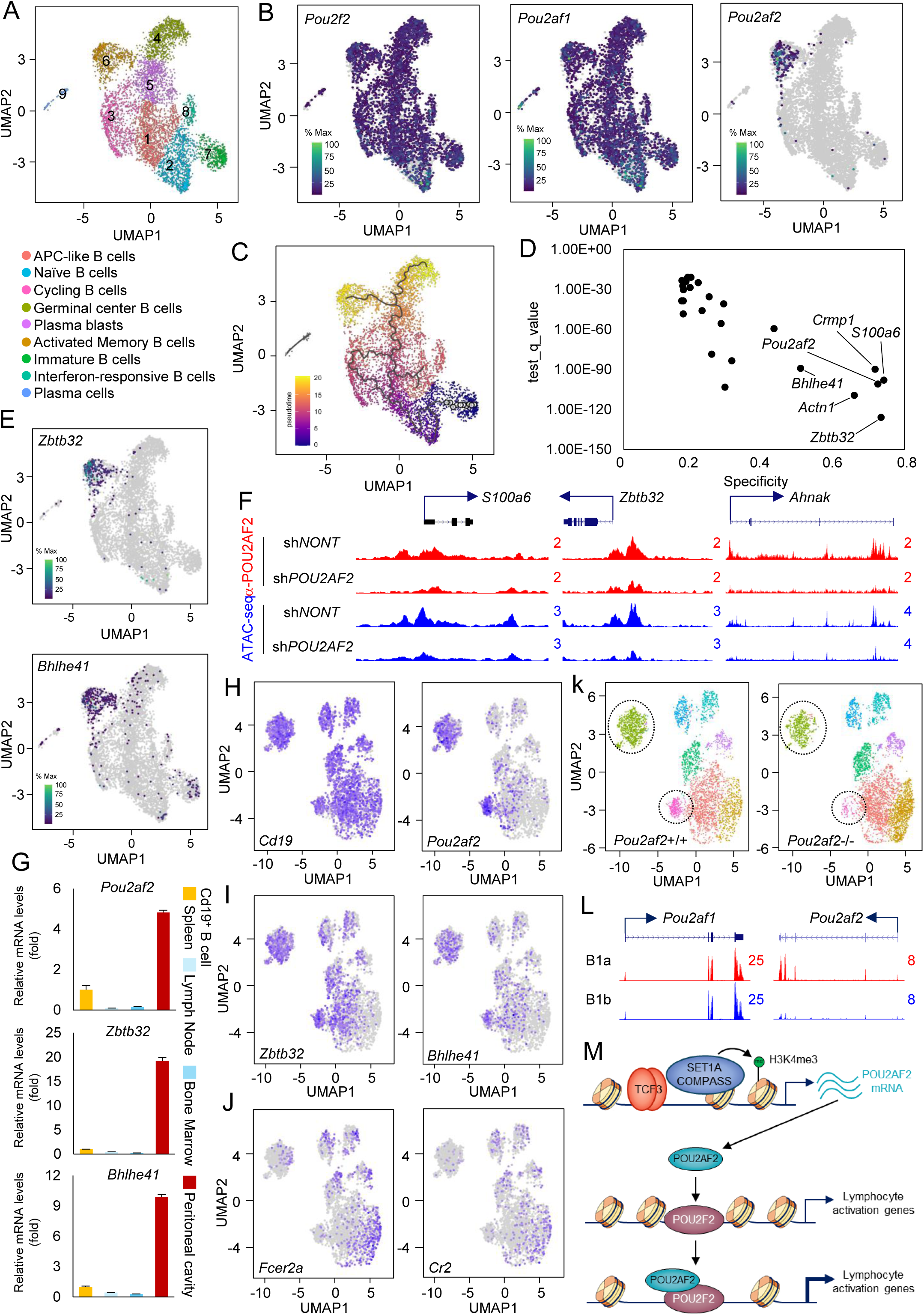
POU2AF2/OCA-T1 defines and sustains a distinct B1-lineage B-cell population *in vivo*. A) Splenic B cells were purified from BALB/c mice and subjected to single-cell RNA sequencing (scRNA-seq). Uniform manifold approximation and projection (UMAP) analysis identifies distinct B cell populations based on transcriptional profiles. B) UMAP plots show cells expressing *Pou2f2* (left), *Pou2af1* (middle), and *Pou2af2* (right). C) Monocle3-based pseudotime analysis of splenic B cells reveals a developmental trajectory of B cell differentiation. D) Scatter plot shows the genes significantly enriched in cluster 6 (*Pou2af2*⁺) cells compared with all other clusters. E) UMAP plots highlights the cell populations expressing *Zbtb32* (top) and *Bhlhe41* (bottom). F) Representative tracks show POU2AF2 occupancy at the genomic loci of genes enriched in cluster 6 and corresponding changes in chromatin accessibility upon POU2AF2 depletion in CH1 cells. G) CD19⁺ B cells were isolated from spleen, lymph node, bone marrow, and peritoneal cavity, and *Pou2af2* mRNA levels were quantified by RT–qPCR, n = 3). H-J) Total peritoneal cavity cells from age- and sex-matched *Pou2af2* wild-type and knockout mice were subjected to scRNA-seq. CD19⁺ B cells were identified by UMAP analysis, and expression of *Cd19* and *Pou2af2* (H), *Zbtb32* and *Bhlhe41* (I), and *Fcer2a* and *Cr2* (J) is shown. K) UMAP analysis of peritoneal cavity B cells from *Pou2af2* wild-type (left) and knockout (right) mice reveals altered B cell cluster distributions. L) Representative genome tracks show expression of *Pou2af1* and *Pou2af2* genes in B1a (CD19^+^B220^lo^CD5^hi^IgM^+^CD23^-^) and B1b B cells (CD19^+^B220^lo^CD5^-^IgM^+^CD23^-^) isolated from mice peritoneal cavity (GSE93764). M) Model of TCF3/SET1A axis–mediated activation of POU2AF2 and its role in promoting lymphocyte activation through co-activation of POU2F2 in B-cell lymphoma.

Next, to identify the genes co-enriched with *Pou2af2* within the same B-cell cluster, we compared gene expression profiles across individual B-cell populations. This analysis identified 131 genes that are preferentially co-expressed with *Pou2af2* gene in the specific B cell population (Fig. 6D), including *Zbtb32*, *Actn1*, *Crmp1*, *S100a6*, and *Bhlhe41* (Fig. 6E, S6B, C). Notably, several of these genes are direct transcriptional targets of POU2AF2 in CH1 cells, and loss of POU2AF2 resulted in a marked reduction in chromatin accessibility at their corresponding genomic loci (Fig. 6F). This result indicates that POU2AF2 may have a similar role in the context of normal B cells, compared to B cell lymphoma cells.

As the two genes most strongly co-expressed with *Pou2af2* gene, *Zbtb32* and *Bhlhe41* are well-established regulators of B-cell differentiation and function(*51–53*). Interestingly, both factors are highly upregulated in B1 B cells, a distinct innate-like B-cell subset that provides rapid, first-line immune defense through the production of natural antibodies against common microbial and self-antigens(*54–56*). B1 B cells are predominantly enriched in the pleural and peritoneal cavities, with smaller populations present in the bone marrow, spleen, and lymph nodes(*57*). We therefore sought to determine the distribution of *Pou2af2*⁺ B cells across immune compartments. As shown in Fig. 6G, *Pou2af2* gene expression was markedly enriched in CD19⁺ B cells isolated from the peritoneal cavity compared with those from lymph nodes, spleen, and bone marrow.

Given that POU2AF2 is required for the viability of both tuft cell-like tumors and normal tuft cells, we next asked whether POU2AF2 similarly regulates the survival of B cells derived from the peritoneal cavity. To test this, total cells were isolated from the peritoneal cavities of sex- and age-matched *Pou2af2* wild-type and knockout mice and subjected to single-cell RNA sequencing. Unsupervised clustering revealed distinct immune populations, including macrophages (*Adgre1*⁺), T cells (*Cd3d*⁺), and B cells (*Cd19*⁺) (Fig. S6D). Consistent with expectations, the majority of *Pou2af2*⁺ cells were Cd19⁺, reflecting the predominance of B cells in the peritoneal cavity (Fig. S6E). Notably, although most Cd19⁺ cells expressed high levels of *Pou2f2* and *Pou2af1* (Fig. S6F), *Pou2af2* expression was largely restricted to B-cell populations enriched with the B1 B cell markers *Zbtb32* and *Bhlhe41* and seen to be depleted of the B2 B cell markers *Fcer2a* and *Cr2* (Fig. 6J). Strikingly, genetic ablation of *Pou2af2* led to a pronounced reduction of this *Pou2af2*⁺ B-cell population in the peritoneal cavity (Fig. 6K), indicating that POU2AF2 is also required for the survival of this subset of normal B cells, which most likely corresponds to the B1 B-cell compartment.

Finally, we analyzed published RNA-seq datasets derived from peritoneal cavity B-1a (*Cd19*^+^*B220*^low^*Cd5*^+^) and B-1b (*Cd19*^+^*B220*^low^*Cd5*^-^*IgM*^+^*Cd23*^-^) cells to compare *Pou2af2* expression between these two cell populations(*51*). As shown in Figure 6L, *Pou2af2* expression was markedly elevated in B-1a cells when compared to B1b cells, whereas *Pou2af1* expression did not differ significantly between the two subsets. Consistent with this observation, *Cd5* expression (a defining marker of B1a cells) was enriched in the *Pou2af2*⁺ B-cell population in both the peritoneal cavity (Fig. S6G) and the spleen (Fig. S6H) in our system. Together, our study defines a cell type–specific epigenetic network that drives POU2AF2 expression, revealing it as a previously unrecognized co-activator of POU2F2 that orchestrates lymphocyte activation programs and sustains both normal B-cell function and B-cell lymphomagenesis (Fig. 6K).

## Discussion

POU2AF2 was initially identified as a novel co-activator of POU2F3, a member of the POU family of transcription factors in tuft cells(*20, 31, 32*). POU2F3 acts as a master regulator that defines a distinct non-neuroendocrine subtype of small-cell lung cancer, which is thought to arise from pulmonary tuft cells within the tracheobronchial epithelium(*22*). In SCLC-P cells, POU2AF2 could be recruited to chromatin by POU2F3, where it bounds to POU2F3 in complex with DNA to enhance the transcriptional activity of POU2F3(*31*). Additionally, we and others have demonstrated that POU2AF2 is able to recruit epigenetic machinery, including the SWI/SNF (BAF) chromatin-remodeling complex, thereby modulating genome architecture that is essential for sustaining the expression of tuft-cell signature genes(*39, 58, 59*). As a key dependency in this cancer subtype, loss of POU2AF2 results in upregulation of tumor suppressors such as PTEN and induces profound apoptosis, leading to near-complete abrogation of tumor growth *in vivo*(*39*).

In the current studies, we have identified a distinct cell type beyond lung epithelial cells, the B cell lymphoma, that expresses high levels of POU2AF2 in both cell lines and primary patient samples. At the chromatin level, POU2AF2 forms a heterodimer with POU2F2, the predominantly expressed POU2 family transcription factor in B cells, to sustain chromatin accessibility and transcriptional output. Notably, in SCLC cells, the vast majority of POU2AF2 occupancy occurs at distal enhancer elements, where the POU2AF2/POU2F3 heterodimer is essential for regulating cell identity genes and cellular plasticity(*39*). By contrast, in the context of B cell lineage, more than two-thirds of POU2AF2 binding sites are localized to promoter regions, and loss of POU2AF2 selectively impairs the expression of functional gene programs, including those involved in lymphocyte activation. Together, these findings reveal the context-dependent transcriptional programs driven by the same co-factor across distinct cancer types, likely reflecting the divergent regulatory functions of POU2 family transcription factors.

TCF3 (E2A) is a transcription factor critical in the differentiation process of common lymphoid progenitors into B-lineage cells and also serves as a central oncogenic driver in B-cell lymphomagenesis(*60–64*). In normal B-cell development, TCF3 orchestrates lineage commitment and maturation by regulating core transcriptional programs that control V(D)J recombination, cell survival, and proliferation(*65*). In B-cell lymphomas, however, dysregulated TCF3 activity, such as recurrent gain-of-function mutations, drives constitutive expression of pro-survival and proliferative genes, including components of B-cell receptor and PI3K signaling pathways(*66*). This aberrant transcriptional output enhances tonic signaling, promotes resistance to apoptosis, and supports malignant transformation and maintenance(*66*). Here, we demonstrate that TCF3 acts as a direct transcription factor driving POU2AF2 expression across multiple B-cell lymphoma cell lines. Notably, although TCF3 is highly expressed in the B-cell lineage, its expression is not restricted to B cells. For example, SCLC cells also exhibit detectable TCF3 protein; however, TCF3 does not constitute a functional dependency in this context. These observations highlight a context-dependent function of TCF3 and warrant future investigation into its chromatin-associated regulatory mechanisms across distinct cellular lineages.

As a major histone H3 lysine 4 (H3K4) methyltransferase, the SET1A COMPASS complex is required to maintain H3K4me3 levels at the promoter of the *Pou2af2* gene. Depletion of SET1A led to a marked reduction in both H3K4me3 enrichment and *Pou2af2* transcription. Notably, pharmacological restoration of H3K4me3 with KDM5 inhibitor in SET1A-depleted CH1 cells failed to rescue *Pou2af2* expression, indicating that H3K4me3 enrichment alone is insufficient to drive transcriptional activation. These findings suggest that SET1A regulates *Pou2af2* expression through mechanisms beyond H3K4me3 deposition. Therefore, our studies highlight therapeutic targeting of SET1A stability by small molecule degraders and the genetic interaction between SET1A COMPASS and TCF3 by small molecule inhibitors as a promising strategy for B-cell lymphoma treatment (Fig. 6k).

POU2AF2 was originally identified as a cell type-specific transcriptional regulator in tuft cells and tuft cell-like tumors, and its genetic ablation results in a global loss of tuft cells in mice(*20, 31, 32*). However, the expression and function of POU2AF2 in other cell types remain poorly understood. Although POU2AF2 mRNA and protein are largely undetectable in bulk mouse splenic B cells, we hypothesized that its expression may be restricted to a distinct B-cell subset. Indeed, single-cell RNA-seq analysis revealed that the vast majority of mouse splenic B cells express *Pou2af1*, the predominant co-activator of POU2F1/2 that is essential for B-cell differentiation and antibody repertoire selection(*67, 68*). Notably, we identified a distinct population of B cells in the normal mouse spleen that express Pou2af2 and are distinguished by elevated expression of B1 B-cell markers, including Zbtb32 and Bhlhe41, relative to other B-cell populations. Consistent with these findings, *Pou2af2*-positive B cells were markedly enriched in B cells derived from the peritoneal cavity, which is known to harbor the highest abundance of B1 B cells. Furthermore, paralleling our observations in B-cell lymphoma, genetic depletion of POU2AF2 significantly reduced the viability of this specific peritoneal B-cell population, suggesting a potential role for POU2AF2 in B1 B cell-mediated rapid innate immune responses and natural antibody production. Taken together, these findings identify POU2AF2 as a context-dependent co-activator regulated by TCF3/SET1A COMPASS axis that integrates transcriptional and epigenetic mechanisms to sustain lineage-specific gene programs in both normal B cells and B-cell lymphoma.

## Methods

### Antibodies and Reagents

POU2F3 (#36135S), Histone H3 (#14269), POU2AF1 (#33483), MLL1 (#14197), MLL2 (#38058S) antibodies were purchased from Cell Signaling Technology. GAPDH (sc-32233) and HSP90 (sc-7947) antibodies were purchased from Santa Cruz. POU2F2 (ab179808) antibody was purchased from Abcam. Flag antibody (F1804) was purchased from Millipore Sigma. TCF3 antibody was purchased from Proteintech (21242-1-AP). SET1A and POU2AF2 antibodies were made in house. R406 (HY-12067) and Ibrutinib (HY-10997) were purchased from (medchemexpress).

### Cell Lines

The HEK293T cells were purchased from ATCC, and maintained with DMEM (Fisher Scientific, #15013CV) containing 10% FBS (Sigma). Cells were adherently cultured in tissue culture-treated flasks at 37°C, 5% CO₂. Media was replaced every 2–3 days and cells were split at ∼70–90% confluency using 0.05% trypsin-EDTA for cell detachment. The A20, 2PK-3, CH1, NCI-H526, NCI-H211, COR-L311 (obtained from ATCC), and the KML-1 cell line (obtained from Japanese Collection of Research Bioresources) were maintained with ATCC-formulated RPMI-1640 medium containing 10% FBS (Sigma). Media was replaced every 2–3 days.

### Mouse experiments

The proposed research activities with vertebrate animals are conducted in the IACUC and AAALAC-approved Center for Comparative Medicine (CCM) facilities. This study was approved by Northwestern University Institutional Animal Care and Use Committee (Animal Protocol No. IS00013610). All animals used in this study were conducted in compliance with the ethical guidelines. Mice were housed five per cage and maintained under specific pathogen-free conditions. Twelve hours of light are provided each day. Food and water were provided freely. 8∼10-week-old BALB/c mice purchased from Jackson lab were used for B cell isolation experiments. 6-week-old nude mice purchased from Jackson lab were used for xenograft experiments. The tumor growth was measured every other day using a calibrated caliper two weeks after inoculation. A two-tailed unpaired Student’s t-test was used for statistical analysis. **P < 0.01; *P < 0.05. Pou2af2-/-mice were generated on a C57Bl6 background, as described(*20*). These mice are maintained in monohybrid cross per Cold Spring Harbor Laboratory IACUC guidelines (Animal Protocol No. 2021-1217). Mice are housed no more than five per cage, with twelve hours of light per day, and food and water provided freely. After birth, tail samples are collected for genomic DNA extraction and genotyping by Sanger sequencing. All the genotyping was performed by Transnetyx, through their automated genotyping platform.

### CRISPR-mediated Knockouts

The mouse sgRNAs were obtained from Mouse CRISPR Knockout Pooled Library (Brie) and human sgRNAs were obtained from Human CRISPR Knockout Pooled Library (Brunello) (PMID: 26780180). The sgRNAs were then cloned into lentiCRISPR v2 (Addgene, 52961) vector. The lentiviral-mediated CRISPR/Cas9 knockout was described previously(*69*). Oligo sequences used in this manuscript were as follows: sg*NONT* (GCTGAAAAAGGAAGGAGTTGA), sg*Tcf3*-1 (GCTCCTAGGAACGTGGAAGG), sg*Tcf3*-2 (GCTGCAAACTGGGTTCCCCCG), sg*Set1a*-1 (GTAGACTGGGCCAAGAGTGG), sg*Set1a*-2 (GCCCCGCACTCGAAAGCACCT), sg*Mll1*-1 (GCTGAGGTGGTATCGATACTG), sg*Mll1*-2 (GCCAAAGCATCGGATCGAGG), sg*Mll2*-1 (GTGAACCCTCTACTCCCCGA), sg*Mll2*-2 (GCCGGGGTGTCTTGATAACAC).

### RNA Interference and Real-time PCR

The cells were infected with lentivirus expressing shRNAs in the presence of 4 μg/ml Polybrene (Sigma) for 24 hours in RPMI-1640 supplemented with 10% FBS. The infected cells were then selected with 2 μg/ml puromycin for an additional 48 hours. The shRNA constructs were purchased from Sigma. The clone IDs for mouse *Tcf3* are TRCN0000233412 (sh*Tcf3*-#1) and TRCN0000233415 (sh*Tcf3*-#2). The clone IDs for mouse *Nfκb2* are TRCN0000012343 (sh*Nfκb2*- #1) and TRCN0000235403 (sh*Nfκb2*-#2). The clone IDs for mouse *Pou2f2* are TRCN0000245326 (sh*Pou2f2*-#1) and TRCN0000432692 (sh*Pou2f2*-#2). The shRNA sequence for mouse *Pou2af2* are ACCCAACTCATTTCGTCTTTA (*shPou2af2*-#1) and TCTTTCATGACAGTGTCAAAT (*shPou2af2*-#2). The non-targeting (sh*NONT*) shRNA construct (SHC002) was purchased from Sigma. Primers for QPCR are as follows, forward: 3′-AACAGCAACTCCCACTCTTC-5′, reverse: 3′-CCTGTTGCTGTAGCCGTATT-5′ (m*Gapdh*); 3′-GGTGTGAGAGTGAAGCACACAG-5′, reverse: 3′-GTAACCTGACATCACTGG-5′ (m*Pou2af2*); forward: 3′-TTTGACCCTAGCCGGACATAC-5′, reverse: 3′-GCATAGGCATTCCGCTCAC-5′ (m*Tcf3*); forward: 3′-TGGCATCCCCGAATATGATGA-5′, reverse: 3′-TGACAGTAGGATAGGTCTTCCG-5′ (m*Nfκb2*); forward: 3′-CTCGCACCTTCAAGCAACG-5′, reverse: 3′-TCGAAGCGGGAAATGGTCG-5′ (m*Pou2f2*); forward: 3′-AGCACTGTGTGATACCCTGAT-5′, reverse: 3′-GGAAGGGCTTATGTCTTCAACC-5′ (m*Zbtb32*); forward: 3′-ATGTGTAAACCCAAAAGGAGCTT-5′, reverse: 3′-TCGGGCAGTAAATCTTTCAGC-5′ (m*Bhlhe41*); forward: 3′-GTCTCCTCTGACTTCAACAGCG-5′, reverse: 3′-ACCACCCTGTTGCTGTAGCCAA-5′ (h*GAPDH*); forward: 3′-GGAGTGAGAGTGAAGCATACAG-5′, reverse: 3′-GTAACCTGACGTGACCGG-5′ (h*POU2AF2*); 3′-ACTGCAGAAGCATCACTCGG-5′, reverse: 3′-CCTGCTTTGTTCTCAGAGCCT-5′ (m*Pou2af2* promoter locus).

#### Immunoprecipitation (IP)

The IP experiment was conducted as previously described(*70*). Briefly, the cells were lysed in the lysis buffer (50mM Tris pH 8.0, 150 mM NaCl, 0.5% Triton X100, 10% Glycerol, protease inhibitors, and benzonase). After centrifugation at 20,000g at 4°C for 15 min, the supernatants were collected and incubated with the antibody and immobilized Protein A/G (Santa Cruz, sc-2003) at 4°C overnight with rotation. Then the Protein A/G beads were washed with ice-cold lysis buffer four times and boiled in 5× SDS sample loading buffer.

### RNA-seq and analysis

RNA-seq was conducted as previously described(*70*). All the steps for library construction were used according to the manufacturer’s recommendations. The RNA-seq samples were pooled and sequenced on a HiSeq with a read length configuration of 150 PE. Gene counts were computed by HTSeq and used as an input for edgeR 3.0.8 Genes with Benjamini-Hochburg adjusted p-values less than 0.01 were considered to be differentially expressed.

### ChIP-seq Assay and analysis

ChIP-seq was performed as described previously(*70*). Briefly, cell pellets were collected, washed twice with ice-cold PBS, and fixed with 1% paraformaldehyde for 10 min at room temperature. Crosslinking was quenched with 2.5 M glycine, followed by two washes with ice-cold PBS. Cells pellets were then resuspended in lysis buffer (50 mM HEPES, pH 7.5, 140 mM NaCl, 1 mM EDTA, 10% glycerol, 0.5% NP-40, 0.25% Triton X-100, and 1× protease inhibitors) and incubated for 10 min at 4 °C with rotation. Nuclei were pelleted by centrifugation at 500 g for 5 min, the supernatant was discarded, and pellets were washed with wash buffer (10 mM Tris-HCl, pH 8.0, 200 mM NaCl, 1 mM EDTA, 0.5 mM EGTA, and 1× protease inhibitors). Washed nuclei were resuspended in sonication buffer (10 mM Tris-HCl, pH 8.0, 1 mM EDTA, 0.1% SDS, and 1× protease inhibitors) and sonicated in 1-ml Covaris tubes using the following settings: 10% duty factor, 175 peak incident power, and 200 cycles per burst for 120–600 s. After sonication, lysates were diluted with 10×dilution buffer (10% Triton X-100, 1 M NaCl, 1% sodium deoxycholate, 5% N-lauroylsarcosine, and 5 mM EGTA) and clarified by centrifugation at maximum speed for 15 min at 4 °C. For immunoprecipitation, approximately 5 μg of commercial antibody or 40 μl of homemade anti-serum was added to each sample and incubated overnight at 4 °C. Protein A/G agarose beads (100 μl) were then added and incubated for an additional 4 h. Beads were then washed four times with RIPA buffer (50 mM HEPES, pH 7.5, 500 mM LiCl, 1 mM EDTA, 1.0% NP-40, and 0.7% sodium deoxycholate), followed by a final wash with ice-cold TE buffer containing 50 mM NaCl. Finally, the DNA was eluted in elution buffer (50 mM Tris-HCl, pH 8.0, 10 mM EDTA, and 1.0% SDS) and reverse cross-linked at 65 °C overnight, followed by proteinase K digestion at 55 °C for 2 h. Genomic DNA fragments were purified using a Qiagen DNA purification kit (Cat. No. 28104). ChIP-seq peaks were called using MACS v2.1.0 with default parameters and corresponding input controls. Metaplots and heatmaps were generated using ngsplot to visualize ChIP-seq signal enrichment. Peak annotation, motif analysis, and super-enhancer identification were performed using HOMER and ChIPseeker, and pathway analysis was conducted with Metascape.

### ATAC-seq and analysis

ATAC-seq was performed as described previously(*71*). Briefly, the frozen cells were thawed, washed once with PBS, and resuspended in ice-cold ATAC lysis buffer. Cell numbers were determined using a Cellometer Auto 2000 (Nexcelom Bioscience). For each sample, 50,000– 100,000 nuclei were pelleted by centrifugation at 500 g for 10 min at 4 °C. After removal of the supernatant, nuclei were resuspended in 50 μl of tagmentation reaction mix and incubated at 37 °C for 30 min in a thermomixer with shaking at 1,000 rpm. Following tagmentation, DNA was purified using the MiniElute Reaction Cleanup Kit (Qiagen) and amplified by PCR using indexed barcode primers. Library quality and concentration were assessed using the Qubit 2.0 DNA High Sensitivity Assay (Thermo Fisher Scientific), the TapeStation High Sensitivity D1000 assay (Agilent Technologies), and the QuantStudio 5 Real-Time PCR System (Applied Biosystems). Libraries were pooled at equimolar concentrations based on quality-control metrics and sequenced on an Illumina HiSeq platform using a paired-end 150-bp configuration, generating approximately 50 million paired-end reads (25 million reads per direction) per sample. For data processing, ATAC-seq reads were shifted by +4 bp and −5 bp for the positive and negative strands, respectively, using the *alignmentSieve* function from the deepTools package. Peaks were identified using MACS v2.1.0.

### Single cell RNA-seq

Cells were washed and resuspended in PBS supplemented with 0.04% BSA. Cell concentration and viability were assessed using acridine orange and propidium iodide staining on a Cellometer Auto 2000 (Nexcelom Bioscience). Dead cells were removed using the Dead Cell Removal Kit (Miltenyi Biotec). Approximately 20,000 viable cells (viability >70%) were loaded onto a Chromium Controller (10x Genomics) following the manufacturer’s protocol for the Chromium 5′ Single Cell Reagent Kit. Following cell partitioning and gel bead–in–emulsion (GEM) generation, reverse transcription was performed, and cDNA was pooled and purified using magnetic beads. cDNA was amplified for 14 PCR cycles and further cleaned using SPRI beads (Beckman Coulter). cDNA quality was assessed using a High Sensitivity D5000 TapeStation assay (Agilent Technologies), and concentration was determined using the Qubit 2.0 DNA High Sensitivity Assay (Thermo Fisher Scientific). 5′ gene expression libraries were prepared according to the Chromium 5′ Single Cell Reagent Kit protocol (10×Genomics). Libraries were pooled at equimolar concentrations based on quality-control metrics and sequenced on an Illumina NovaSeq platform using a paired-end 150-bp configuration, generating approximately 500 million paired-end reads per sample (250 million reads per direction).

### Statistical Analyses

Statistical analyses were performed using GraphPad Prism 7, Microsoft Excel, and R. All datasets met the assumptions required for the corresponding statistical tests. Data distribution was assessed using the empirical rule to evaluate normality. For growth curves and time-course RNA-seq analyses, statistical comparisons were performed using area-under-the-curve (AUC) values. The specific statistical tests applied for each analysis are indicated in the figure legends.

## Data availability

NGS data generated for this study will be provided upon request.

## Acknowledgement

We are grateful for all the past Wang lab members for their support. L.W. is supported by NIH grant R35GM146979, the Research Scholar Grant (RSG-22-039-01-DMC) from the American Cancer Society, and the Idea development award (HT94252310360) from USAMRAA. C.R.V. is supported by the National Institutes of Health R01 grant CA290004.

---

**Supplementary Figure 1.**
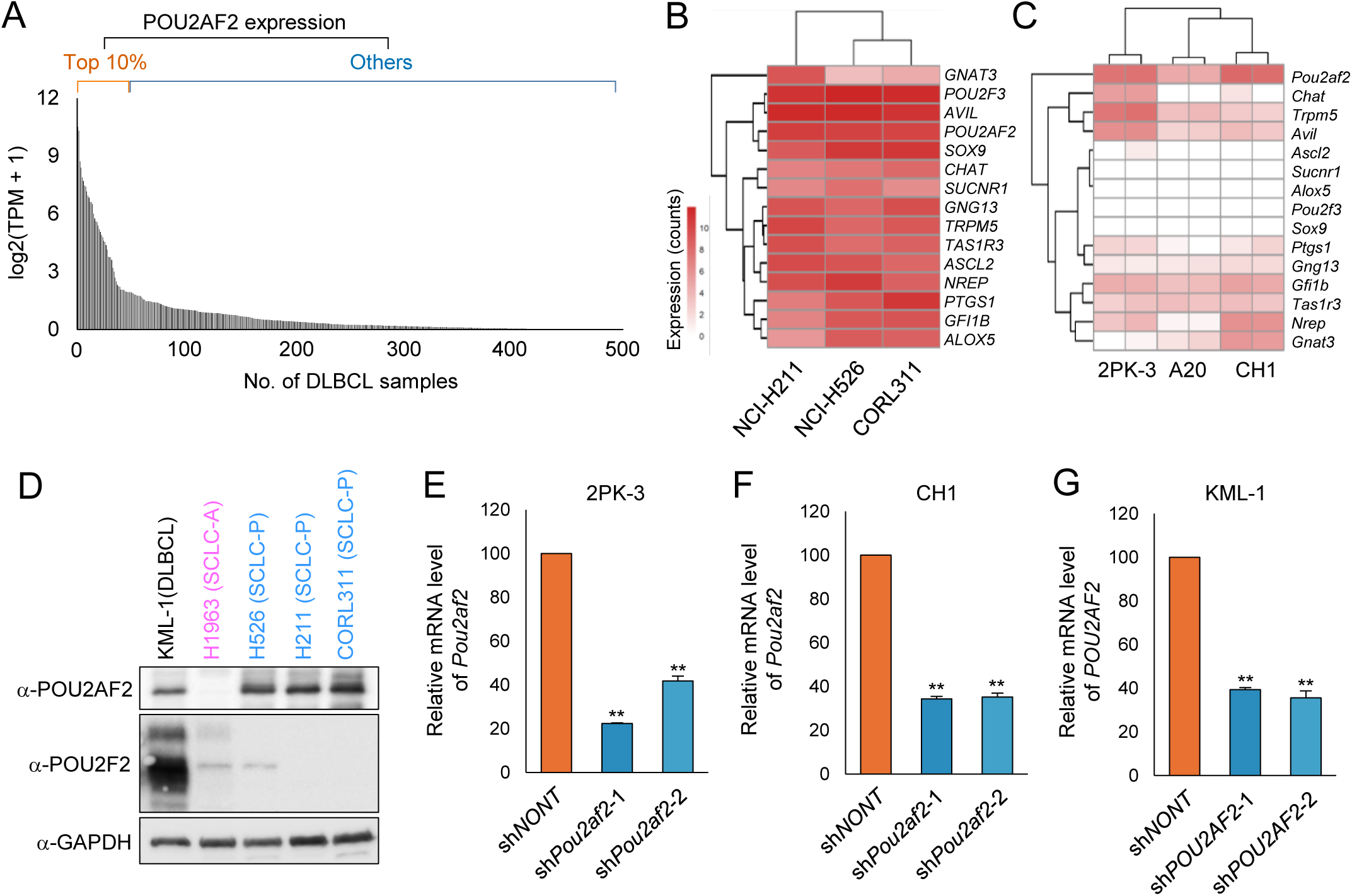
POU2AF2/OCA-T1 is a lineage-specific dependency in a subset of diffuse large B-cell lymphoma. A-B) Expression of tuft cell-signature genes in P-subtype small cell lung cancer (SCLC-P) cell lines (A) and B cell lymphoma cell lines (B). C) Western blot analysis of POU2AF2 and POU2F2 protein levels in the human B cell lymphoma cell line KML-1 and SCLC cell lines NCI-H526, NCI-H211, and CORL311, n=2. D-F) Mouse B cell lymphoma cell lines 2PK-3 (D), CH1 (E), and human B cell lymphoma cell line KML-1 (F) were transduced with nontargeting or two independent *Pou2af2* shRNAs. The *Pou2af2* mRNA levels were quantified by RT–qPCR, n = 3. Data are represented as mean ± SD. A two-tailed unpaired Student’s t-test was used for statistical analysis. **P < 0.01; *P < 0.05.

**Supplementary Figure 2.**
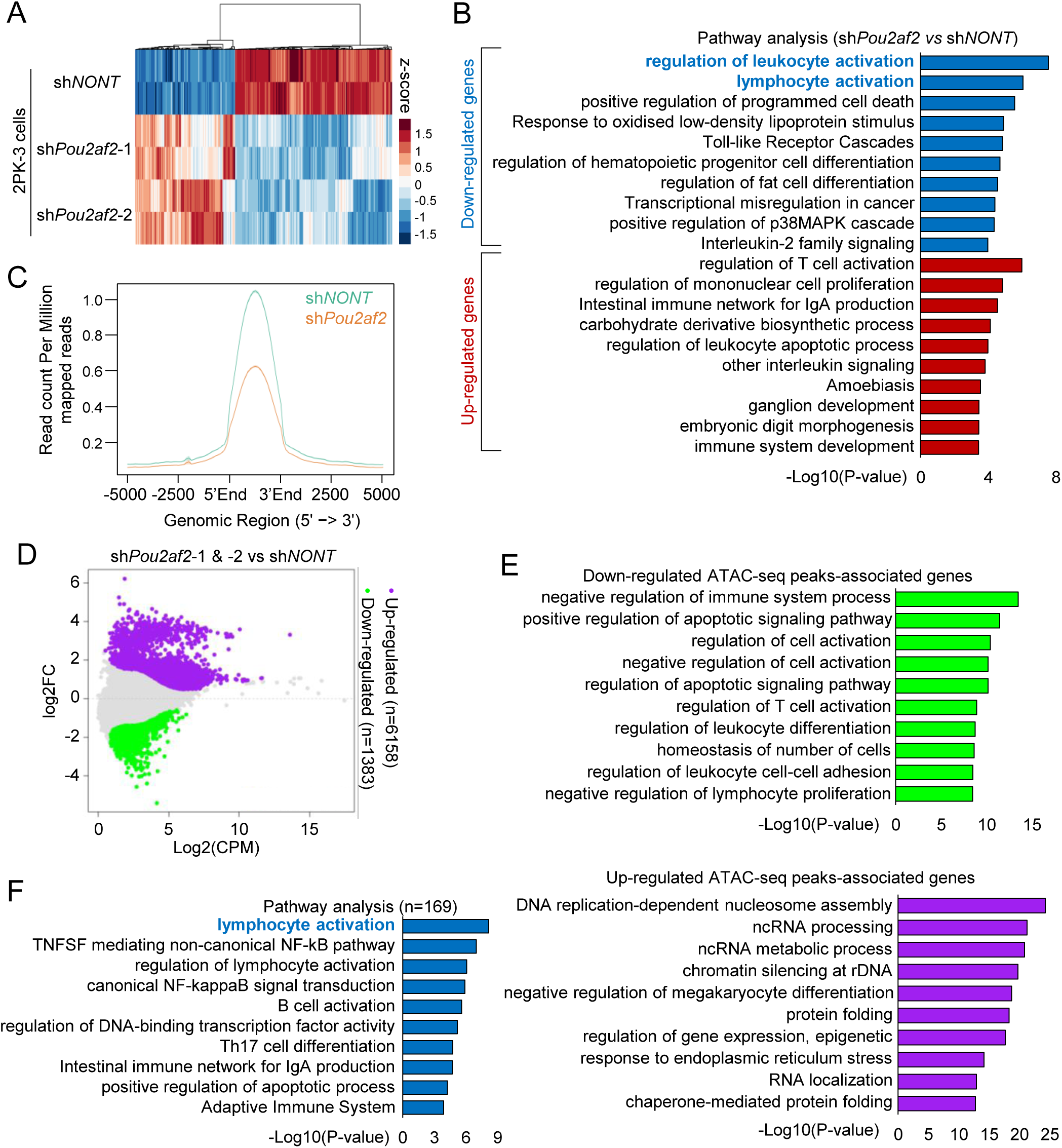
POU2AF2/OCA-T1 controls the expression of lymphocyte activation genes at chromatin. A) RNA-seq analysis of 2PK-3 cells transduced with nontargeting or two independent *Pou2af2* shRNAs for 4 days. Heatmap shows differentially expressed genes, n = 2. B) Metascape pathway enrichment analysis of genes that were upregulated or downregulated upon *Pou2af2* gene depletion. Enrichment significance is shown as −log_10_(P) (Metascape v3.5). C) CH1 cells transduced with nontargeting or *Pou2af2*-specific shRNA were subjected to ChIP–seq with POU2AF2 specific antibody. Average plot shows antibody specificity. D) MA plot shows the significantly upregulated and downregulated ATAC-seq peaks upon *Pou2af2* gene depletion in CH1 cells. E) Metascape pathway enrichment analysis of genes associated with differentially accessible ATAC-seq peaks in *Pou2af2*-depleted cells. F) Pathway analysis of the genes downregulated upon POU2AF2 depletion in CH1 cells and upregulated upon POU2AF2 overexpression in A20 cells (n=169).

**Supplementary Figure 3.**
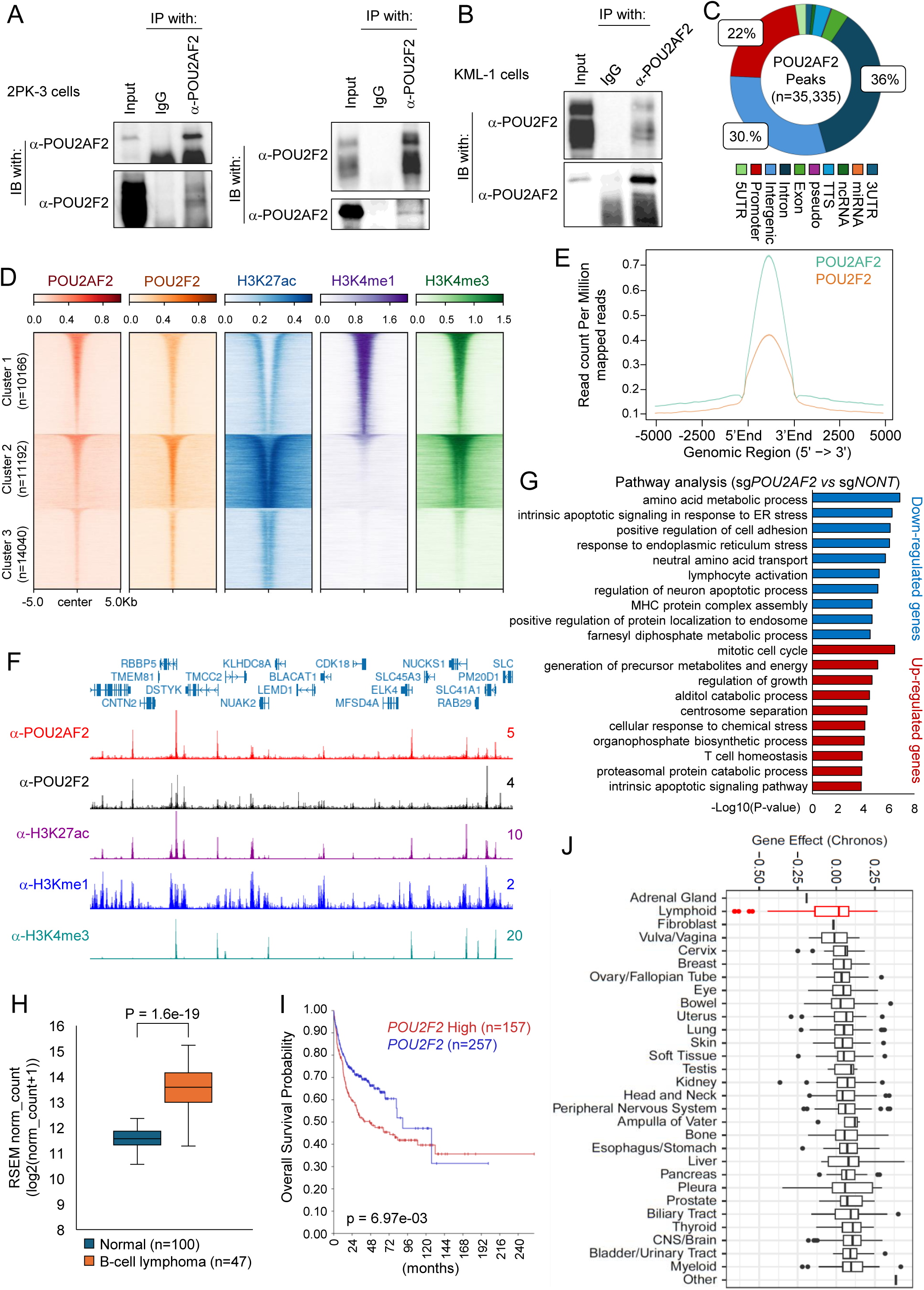
POU2AF2/OCA-T1 co-activates POU2F2 in B cell lymphoma. A-B) Co-immunoprecipitation shows interaction between POU2AF2 and POU2F2 in whole-cell lysates from 2PK-3 (A) and KML-1 (B) cells, n=2. C) Pie chart shows genomic distribution of POU2AF2 ChIP–seq peaks in KML-1 cells. D) The heatmap shows the co-occupancy of POU2AF2 and POU2F2 in KML-1 cells. Rows are centered on three POU2AF2 peak clusters defined by k-means clustering, with histone marks aligned to peak centers. E) Genome-wide average plot shows the co-occupancy of POU2AF2 and POU2F2 in KML-1 cells. F) Representative tracks show co-localization of POU2AF2 and POU2F2. G) KML-1 cells transduced with nontargeting or two independent *POU2AF2* sgRNAs were analyzed by RNA-seq, followed by Metascape pathway enrichment analysis of differentially expressed genes. H) The expression levels of *POU2F2* in normal B cells and B cell lymphoma were retrieved from UCSC Xena database(*35*). The box plot shows *POU2F2* gene expression in normal samples versus human B cell lymphoma. I) Kaplan–Meier analysis by the R2 Genomics Analysis and Visualization Platform (https://hgserver1.amc.nl/) shows the correlation between *POU2F2* expression and overall survival in B cell lymphoma patients. J) Gene dependency scores of POU2F2 across different tumor lineages obtained from the DepMap database.

**Supplementary Figure 4.**
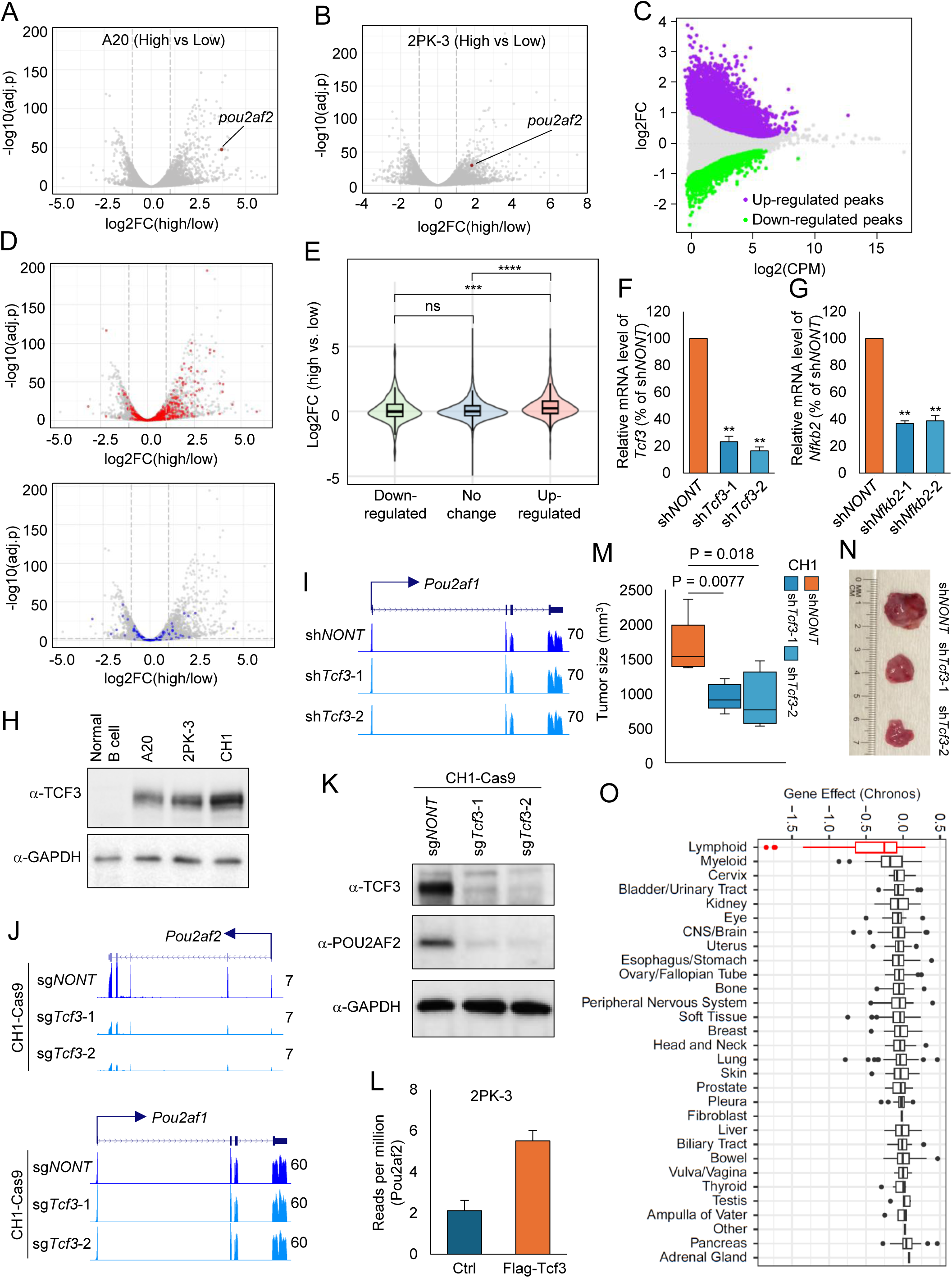
TCF3 links BCR signaling to *POU2AF2* activation and B-cell lymphoma progression. A-B) Volcano plots show differentially expressed genes in A20 (A) and 2PK-3 (B) cells cultured under low- or high-density conditions for 4 days. C) MA plot shows differentially accessible ATAC-seq peaks in A20 cells cultured under low- or high-density conditions. D-E) Volcano (D) and violin (E) plots show the expression of genes associated with differentially accessible ATAC-seq peaks. F-G) RT-qPCR analysis of *Tcf3* (F) and *Nfκb2* (G) mRNA levels in A20 cells transduced with nontargeting or gene-specific shRNAs, n = 3. Data are represented as mean ± SD. A two-tailed unpaired Student’s t-test was used for statistical analysis. **P < 0.01; *P < 0.05. H) Western blot analysis of TCF3 protein levels in mouse splenic B cells, B cell lymphoma cell line A20, 2PK- 3, and CH1 cells, n=2. I) Representative RNA-seq tracks show *Pou2af1* mRNA levels in CH1 cells transduced wither non-targeting shRNA or two independent Tcf3 shRNAs, n=2. J) Representative RNA-seq tracks show *Pou2af2* and *Pou2af1* mRNA levels in CH1-Cas9 cells transduced with non-targeting sgRNA or two independent Tcf3 sgRNAs, n=2. K) Western blot analysis of TCF3 and POU2AF2 protein levels in CH1-Cas9 cells transduced with non-targeting sgRNA or two independent Tcf3 sgRNAs, n=2. L) Bar plot shows the *Pou2af2* gene expression in 2PK-3 cells expressing control vector or FLAG-TCF3, n=2. O) Gene dependency scores of *TCF3* across different tumor lineages from the DepMap database.

**Supplementary Figure 5.**
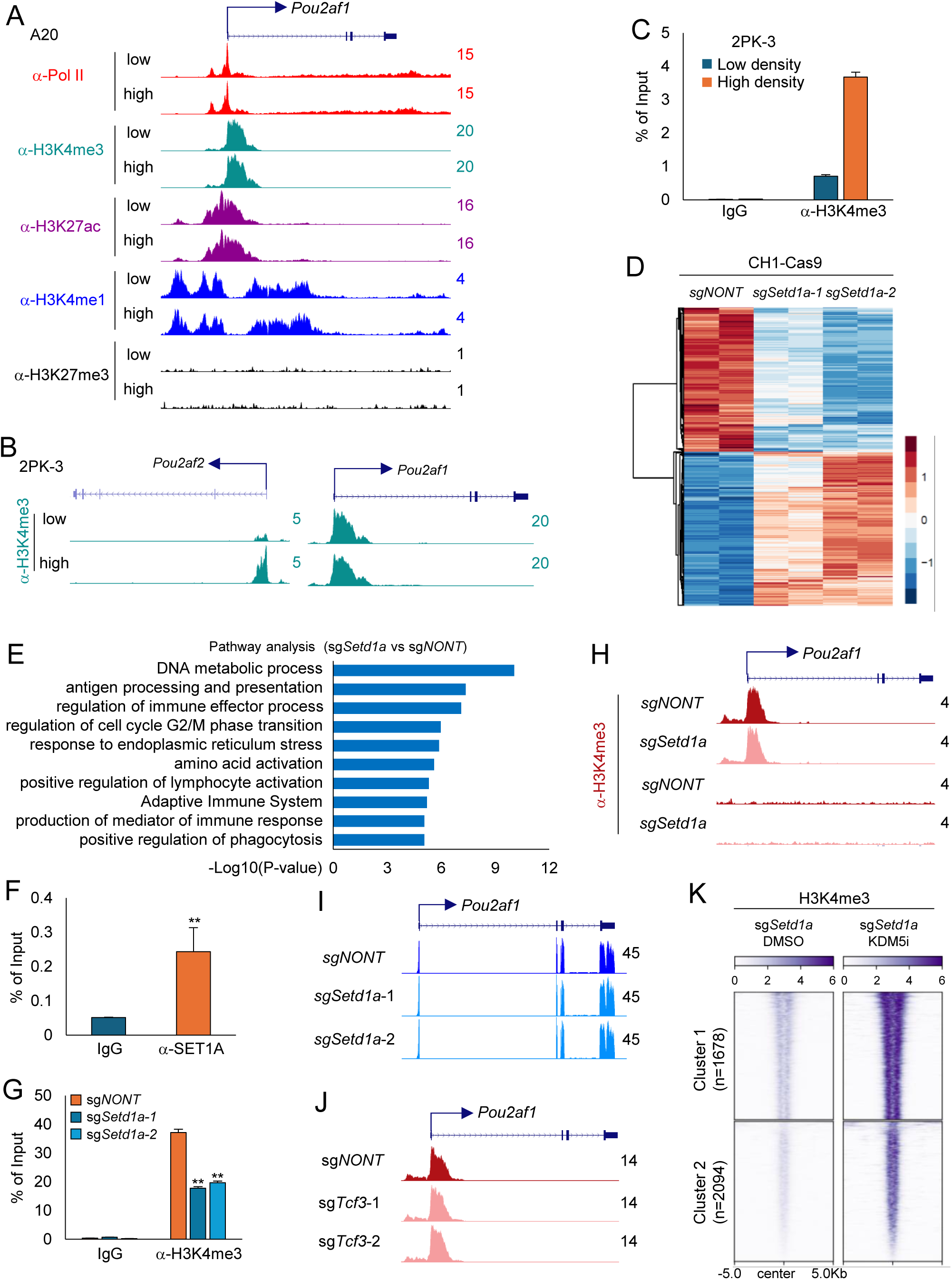
SET1A COMPASS selectively controls *POU2AF2* expression in B-cell lymphoma. A) Representative tracks show Pol II, H3K4me3, H3K4me1, H3K27me3, and H3K27ac occupancy at the *Pou2af1* gene promoter in A20 cells cultured under low- or high-density conditions. B) Representative tracks show H3K4me3 levels at *Pou2af2* and *Pou2af1* gene loci in 2PK-3 cells cultured under low- or high-density conditions. C) ChIP–qPCR quantification of H3K4me3 at the promoter region of *Pou2af2* gene, n=3. Data are represented as mean ± SD. A two-tailed unpaired Student’s t-test was used for statistical analysis. **P < 0.01; *P < 0.05. D) RNA-seq analysis of CH1-Cas9 cells transduced with nontargeting or two independent *Setd1a* sgRNAs. Heatmap shows differentially expressed genes, n=2. E) Metascape pathway enrichment analysis of genes downregulated upon SET1A depletion in CH1-Cas9 cells. F) ChIP-qPCR analysis of SET1A occupancy at the *Pou2af2* promoter in CH1 cells, n=3. Data are represented as mean ± SD. A two-tailed unpaired Student’s t-test was used for statistical analysis. **P < 0.01; *P < 0.05. G) ChIP-qPCR analysis of H3K4me3 levels at the *Pou2af2* promoter in CH1-Cas9 cells transduced with either nontargeting or two independent *Setd1a* sgRNAs, n=3. Data are represented as mean ± SD. A two-tailed unpaired Student’s t-test was used for statistical analysis. **P < 0.01; *P < 0.05. H-I) Representative tracks show H3K4me3 levels (H) at the promoter region of *Pou2af1* gene (I) and *Pou2af1* gene expression in CH1-Cas9 cells transduced with either nontargeting or *Setd1a* sgRNA. J) Representative tracks show H3K4me3 occupancy at the *Pou2af1* gene promoter upon TCF3 depletion in CH1-Cas9 cells. (K) The heatmap shows H3K4me3 ChIP–seq signal in SET1A-depleted CH1 cells treated with DMSO or 5 μM KDM5 inhibitor for 24 hours.

**Supplementary Figure 6.**
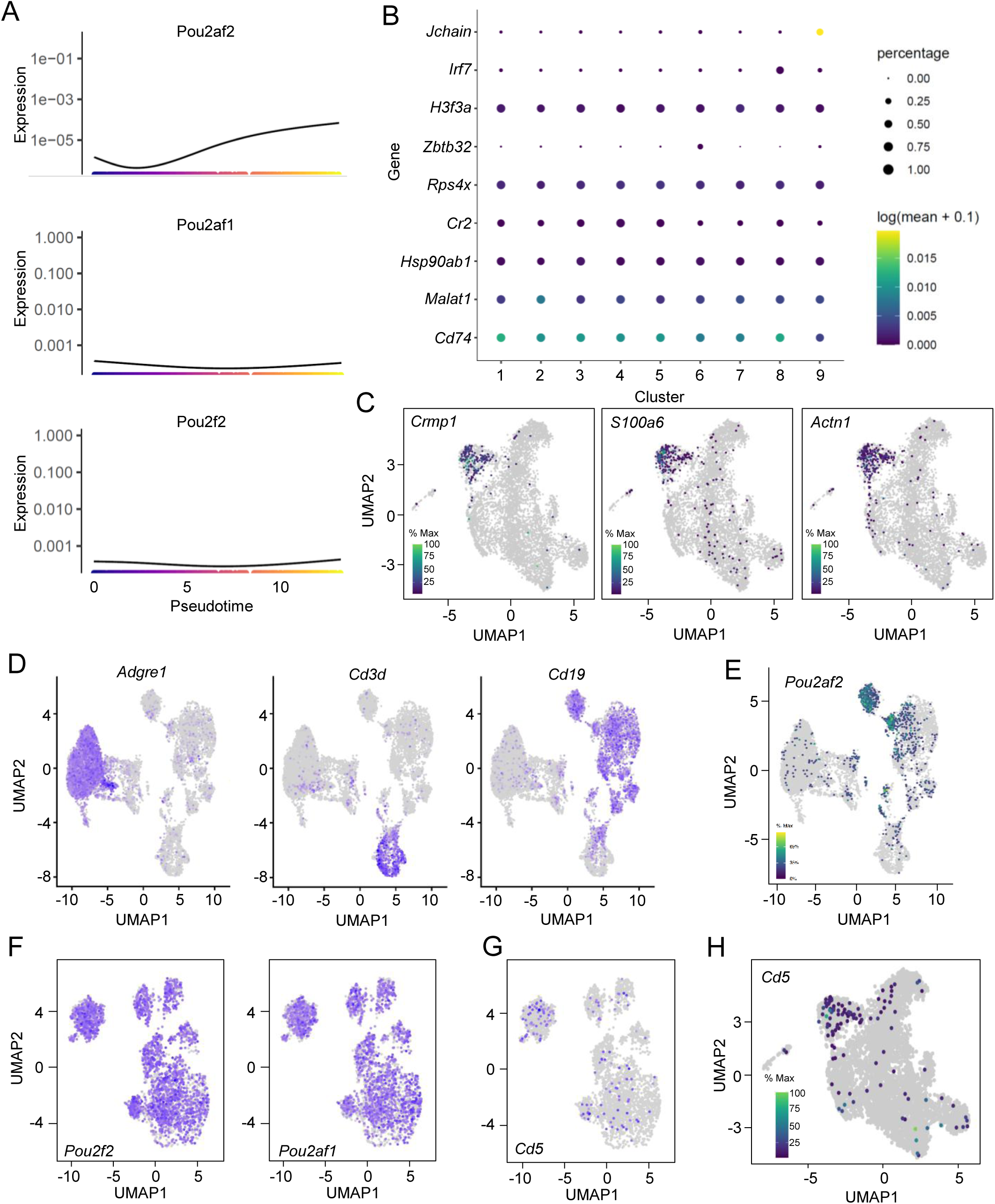
POU2AF2/OCA-T1 defines and sustains a distinct B1-lineage B-cell population *in vivo*. A) Monocle3-based pseudotime analysis showing expression dynamics of *Pou2af2*, *Pou2af1*, and *Pou2f2* gene expression during cell development. B) Dot plot shows the top enriched gene in each UMAP-defined cluster. C) UMAP plots identify Crmp1-, S100a6-, and Actn1-positive cell populations. D) scRNA-seq analysis of total peritoneal cavity cells identify macrophages (*Adgre1*⁺), T cells (*Cd3d*⁺), and B cells (*Cd19*⁺). E) UMAP plot identifies *Pou2af2*-positive cells from total peritoneal cavity derived cells. UMAP analysis of *Cd19*⁺ B cells highlighting *Pou2f2*, *Pou2af1* gene expression (F) and *Cd5*⁺ populations (G). H) UMAP analysis identifies *Cd5*⁺ cells among total splenic B cells.

